# Trans-kingdom coupling of redox signaling to environmental cell stress responses through multiphase partitioning

**DOI:** 10.1101/2025.05.09.653213

**Authors:** Elsy Ngwa, Xi Chen, Stefanie Schmieder, Aaron Maurais, Victor Svistunov, Qianni Peng, Ian Moore, Wayne I. Lencer, Eranthie Weerapana, Krishnan Raghunathan, Jay R. Thiagarajah

**Author notes:** These authors contributed equally (Elsy Ngwa, Xi Chen). These authors contributed equally (Krishnan Raghunathan, Jay R. Thiagarajah). **Classification:** Biophysics and Computational Biology; Cell Biology; Evolution.

## Abstract

Reactive Oxygen Species (ROS) signaling is a conserved biological process with parallel functions in all evolutionary branches of life. Here, we identify Receptor for Activated C Kinase 1 (RACK1) as a conserved redox-regulated hub that integrates ROS signals to coordinate cellular stress responses. Using cysteine reactivity profiling in intestinal epithelial cells, we demonstrate that RACK1 undergoes NOX1-dependent oxidation at multiple residues, with C286 serving as a key regulatory site. Functional studies reveal that RACK1 negatively regulates NFκB signaling through redox-dependent interactions with upstream signaling complexes. Upon stress stimulation, RACK1 dynamically redistributes into membrane-less condensates that act as redox-privileged microenvironments enriched for hydrogen peroxide. We find that oxidized RACK1 condensates are conserved through evolution with analogous stress response behavior in bacteria and yeast indicating a ubiquitous and ancient stress sensor-effector system. Functionally, redox-dependent RACK1 activity links environmental stress to translational control, with oxidation promoting inhibition of protein synthesis. Furthermore, RACK1 mediates responses to diverse pathogen-associated stimuli, including viral and bacterial infection, highlighting its role in epithelial innate immune responses. Collectively, these findings establish RACK1 as a cellular node for redox signaling, operating within condensate-based microdomains to spatially encode oxidative signals and regulate environmental stress pathways in cells.

**Significance Statement:** This study defines a mechanism by which cells achieve specificity in redox signaling through compartmentalization within condensates. By identifying RACK1 as a redox sensor-effector, we reveal an ancient and broadly conserved system linking environmental stress to innate immune signaling and translational control. The discovery of redox-privileged condensates provides a conceptual framework for understanding how transient ROS signals are stabilized and interpreted in epithelial biology, with implications for inflammatory diseases, host–pathogen interactions, and fundamental cellular stress responses.

## Main Text

Life on earth began in anoxic conditions and initial cellular organization and biochemical systems emerged in a reducing environment. (1–3) As oxygen levels increased, early organisms evolved to sense and regulate reactive oxygen species (ROS) formed as byproducts of cellular oxygen utilization. (4) Most organisms co-opted relatively stable ROS species such as hydrogen peroxide as a potential sensor-effector system to regulate major cellular processes including respiration and metabolism as well as signaling pathways through reversible redox modifications of specific amino acids such as cysteines.(5, 6) ROS signaling as a response to environmental and pathological stress is especially relevant in tissues interfacing with the external environment such as barrier epithelial cells.(7, 8) Cellular responses in intestinal epithelial cells are modulated by hydrogen peroxide (H_2_O_2_) produced by NADPH oxidases (NOXs). NOX1, one of the major oxidases present in colonic epithelial cells, functions in the cell and tissue response to injury and infection, with loss-of-function associated with inflammatory bowel disease and over-expression with cancer.(9–11) Previous data showed that H_2_O_2_ produced by NOX1 and transported into cells modulates specific inflammatory responses in colonic epithelial cells.(12) To protect from non-specific or excessive ROS, cells maintain a generally reductive cytosolic environment with rapid removal of oxidizing equivalents.(13) To ensure specificity and veracity, H_2_O_2_ signaling requires cellular spatial and temporal compartmentalization. While the general importance of ROS in both cell physiology and pathology are well appreciated, our understanding of the underlying dynamics and mechanistic modules that enable signaling responses continues to evolve. In this work, we have identified that a highly conserved protein - RACK1-acts as a central redox-regulated hub in the sensing and transmission of environmental stress signals in cells. We show that both the protein and its mechanism of redox signal propagation are conserved through evolution and, in the context of the intestine, modulates the response to an array of stressors including pathogens, by negatively regulating NFκB signaling, ribosomal function and protein translation.

## Results

### Receptor for Activated C Kinase 1 (RACK1) is a redox-regulated protein in native intestinal epithelial cells

Previous data using infection models has suggested that environmental signaling is modulated by ROS in barrier intestinal epithelial cells.(12) Since NOX1 is a major source of the stable ROS species, hydrogen peroxide, in colonic epithelial cells,(14) we confirmed in mouse and human cells that loss of NOX1 function leads to altered downstream signaling in response to a prototypic danger-associated molecule, IL-1β. Either genetic loss (NOX1^-/-^) or chemical inhibition (Figure 1A, C), leads to reduced transcription of IL-8 in response to IL-1β stimulation. We observed a general dependence of downstream signaling on NOX1, across diverse stimuli from damage and pathogen-associated molecules such as IL-1α, TNFα, IL-17A, LPS, Flagellin, and pIC (Figure 1B). Since NOX1-derived ROS can modulate protein function through cysteine oxidation, we applied proteome-wide reactive-cysteine profiling to identify potential sites of cysteine oxidation.(15, 16) Colonoids from NOX1^+/+^ and NOX1^-/-^ mice were generated and stimulated with IL-1β. Proteomes from control and treated groups were labeled with isotopically heavy and light iodoacetamide-alkyne (IA-L/H) probes, respectively (Figure 1D). Mass-spectrometry analysis of IA-tagged peptides afforded heavy:light (H/L) ratios that reflect on cysteine reactivity changes induced upon IL-1β treatment. Oxidation of a cysteine upon IL-1β treatment causes a decrease in cysteine reactivity, which is reflected in Log_2_ H/L values >0. Analysis of cysteines that show increased oxidation in NOX1^+/+^ relative to NOX1^-/-^ proteomes revealed GNB2L1, also known as RACK1, as one of the most highly represented differentially oxidized proteins. (Figure 1E, F, S1A). RACK1 is a member of the tryptophan-aspartate repeat (WD-repeat) family of proteins and adopts a seven-bladed β-propeller structure with cysteine residues spaced throughout the protein structure (Figure 1G).(17) RACK1 is a known to chaperone protein kinase C (PKCβII) and a component of ribosomal 40S subunits.(18–20) Our cysteine-profiling data identified at least 6 cysteines that displayed reduced oxidation with loss of NOX1 function (Figure 1H). Conservation analysis of RACK1 revealed remarkable residue conservation through evolution leading back to early eukaryotic organisms (Figure 1I).

**Figure 1.**
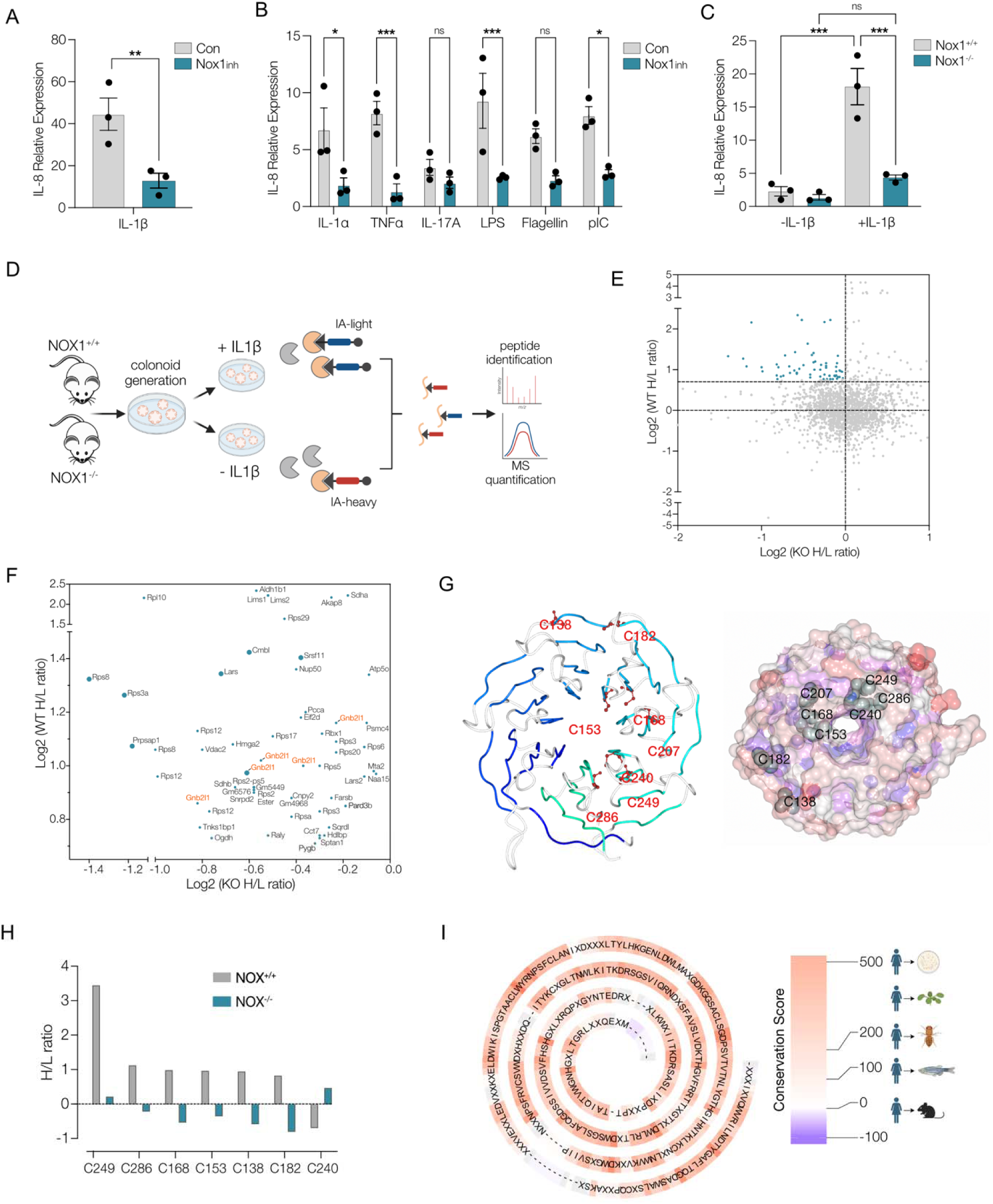
Receptor for Activated C Kinase 1 (RACK1) is a redox-regulated protein in native intestinal epithelial cells. **(A)** Relative expression of IL-8 in Caco2-BBE cells with and without Nox1 inhibitor (ML171, 5μM) in response to IL-1β. (n=3) **(B)** Relative expression of IL-8 in Caco2-BBE cells with and without Nox1 inhibitor (ML171, 5μM) in response to damage and pathogen associated molecules. (n=3) **(C)** Relative expression of IL-8 in NOX1^-/-^ and WT mouse colonoids in response to IL-1β. (n=3) **(D)** Schematic showing the experimental outline for generation of colonoids and isoTOP-ABPP proteomics **(E)** Scatter plot of Log_2_ H/L values in NOX1^-/-^ and WT colonoids with differentially oxidized cysteine residues in WT versus NOX1^-/-^ highlighted in blue. **(F)** Scatter plot of proteins with significantly redox-modified cysteine residues in WT versus NOX1^-/-^ colonoids. Cysteines from GNB2L1 (RACK1) are highlighted in orange. **(G)** Ribbon and space filling model of crystal structure of RACK1 showing the relative positions of cysteine residues. **(H)** H/L ratios of the individual cysteines of RACK1 in both WT and NOX1^-/-^ cells. **(I)** Conservation plot of RACK1 from lower eukaryotes to human. Values represent the conservation score from multiple sequence alignment. Data are presented as mean ± SEM. Statistical significance was determined using two-tailed unpaired Student’s t tests in (A) and (B); ns, not significant; *p < 0.05, **p < 0.01, ***p < 0.001

### Oxidative modification of C286 mediates negative regulation of NF**κ**B signaling

A major cellular effector pathway activated by IL-1β is NF-κB mediated signaling. IL-1β-stimulated NF-κB activity was found to be directly modulated by altering cellular redox status of the cell. As previously established, N-acetylcysteine, which increases cellular glutathione and reduces ROS, decreases NF-κB activity. In contrast, rotenone, a mitochondrial complex I inhibitor which increases ROS in the cell, enhances NF-κB activity (Figure 2A). Overexpression of RACK1 in cells led to a significant decrease in NF-κB activity consistent with previous data on RACK1 function. (Figure 2B) (20) To assess whether redox modification is necessary for RACK1-mediated regulation of NF-κB activity we overexpressed RACK1 with individual or multiple mutations (C to A) of the identified cysteine residues. We found that loss of all redox modifiable cysteines (C-All) led to a significant negative regulation of NF-κB activity (Figure 2C). We observed that equivalent negative regulation was induced by mutating a single specific cysteine, C286, suggesting that this residue is particularly critical for this function. In all subsequent experiments, we used either C286A or C-All as non-reactive and expression controls to WT RACK1. Negative regulation of NF-κB by RACK1 was dose-responsive with activity a log-fold lower at low doses with the C286 mutant versus WT RACK1. (Figure 2D). Conversely knockdown of RACK1 led to increased NF-κB activation (Figure S1B). Activation of NF-κB transcription requires phosphorylation and nuclear translocation of the protein p65. Assessment of nuclear localization of p65 upon IL-1β stimulation showed that increased expression of WT RACK1 reduced nuclear p65 relative to control which was further decreased with C286A-RACK1 (Figure 2E, F). Similarly, p65 phosphorylation was reduced with WT-RACK1 expression and further reduced with C286A-RACK1 indicating negative regulation by RACK1 occurs upstream of p65 (Figure 2G). Previous studies have shown functional interactions between RACK1 and the IKK complex, specifically with IKKα.(20) Co-immunoprecipitation showed that at baseline endogenous RACK1 associates with IKKα but rapidly dissociates following stimulation with IL-1β (Figure 2H). In contrast, IKKα remained associated with overexpressed WT RACK1 following IL-1β, and this association was further increased in cysteine mutants (C-All) indicating that redox-regulation of RACK1 state is involved in inhibition of NFκB activity under tonic and stimulated conditions. We also observed that cysteine-dependency of RACK1 regulation of NFκB activity was not restricted to IL-1β stimulation but conserved for other known pathway activators including TNFα and LPS (Figure 2J).

**Figure 2.**
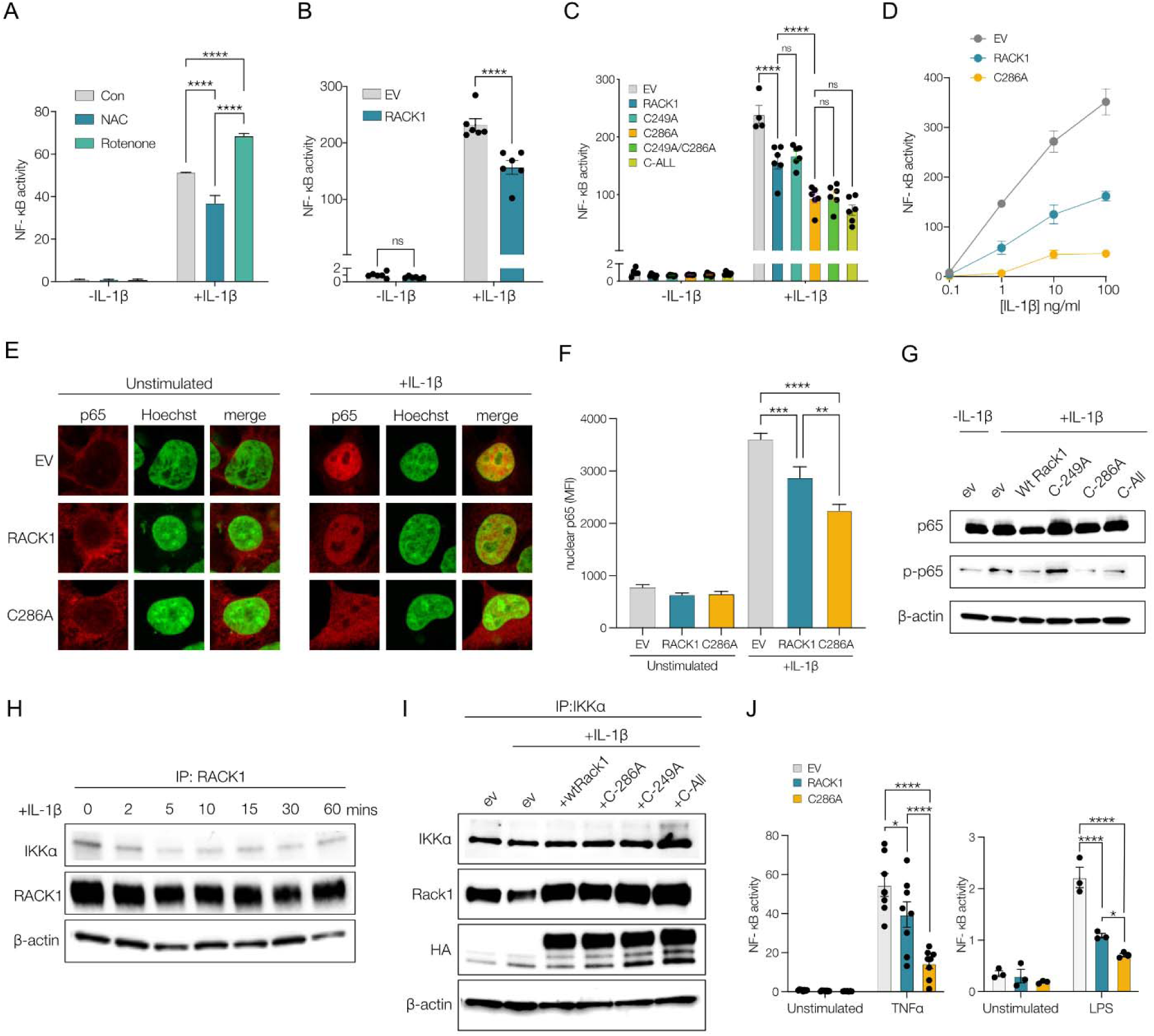
Oxidative modification of C286 mediates negative regulation of NFκB signaling. **(A)** NF-κB activity in HEK cells pretreated with either media, N-acetyl Cysteine (NAC, 5mM) or Rotenone (1mM) and stimulated with IL-1β (10 ng/μL) or control. (n=3) **(B)** NF-κB activity in HEK cells transfected with RACK1-WT, or empty vector (EV) stimulated with IL-1β (10 ng/μL) or control. (n≥4) **(C)** NF-κB activity in HEK cells transfected with RACK1-WT, RACK1 cysteine mutants or empty vector (EV) stimulated with IL-1β (10 ng/ml) or with control. (n≥4) **(D)** NF-κB activity in HEK transfected with RACK1-WT, C286A mutant or empty vector (EV) with varying dose of IL-1β. (n≥4) **(E)** p65 immunofluorescence staining in empty vector, RACK1-WT or RACK1-C286A transfected HEK cells either with mock or IL-1β (10 ng/ml) stimulation for 30 minutes. Nuclei were stained using Hoechst. **(F)** Quantification of immunofluorescence images from (E). (n≥2) **(G)** Western blot graph showing relative p65 phosphorylation in HEK293T expressing RACK1-WT, RACK1 cysteine mutants or empty vector **(H)** Time series of co-immunoprecipitation of IKK〈 with overexpressed wtRACK1 in HEK293T following IL-1β (10 ng/μL) stimulation. **(I)** Co-immunoprecipitation of IKKα with RACK1 in HEK293T expressing empty vector (ev), wt or cysteine mutants RACK1 following IL-1β (10 ng/μL,1h) stimulation. **(J)** NF-κB activity in HEK cells expressing empty vector (ev), wt or cysteine mutants RACK1 stimulated with TNFα, LPS or unstimulated. *p < 0.05, **p < 0.01, ***p < 0.001

### RACK1 forms stress-induced redox-privileged puncta

Given that RACK1 redox modification appeared to be a generalized feature of the cellular signaling response to several stimuli, we investigated the cellular location of RACK1 in stress sensing. HCT-8 cells transfected with RACK1-GFP, were stimulated with either exogenous stress (IL-1β) or endogenous stress (heat-shock protein inhibitor 17-AAG). We observed that upon stimulation, RACK-1 rapidly redistributed from a primarily cytoplasmic pattern to enriched punctate structures that grow in both number and size over time. (Figure 3A, 3B). HCT-8 transfected with RACK1-mKate cells also formed condensates when treated with H_2_O_2_ (Figure S2A). We observed that the intensity of the puncta was highest following 17-AAG treatment and used this condition to measure the properties of RACK1 puncta. Puncta were also observed with endogenous RACK1 (Figure 3C) following 17-AAG treatment.

**Figure 3.**
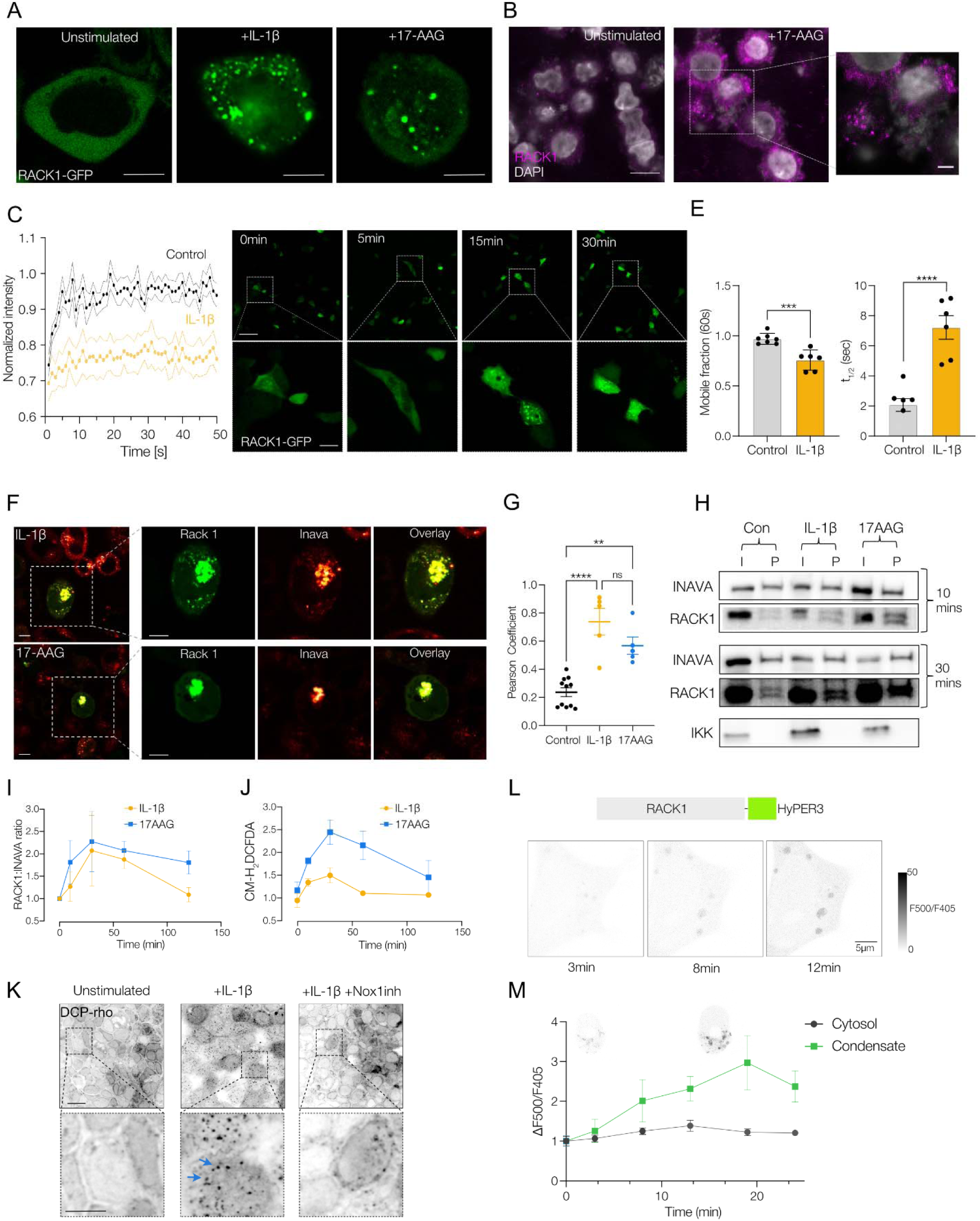
RACK1 forms stress-induced redox-privileged puncta. **(A)** Representative fluorescence image showing RACK1 condensates in HCT-8 cells expressing RACK1-GFP and stimulated with IL-1β (10 ng/μL) or with 17-AAG (10 µM) but not in untreated cells. **(B)** Representative immunofluorescence of HCT-8 showing endogenous RACK1 puncta when treated with 17-AAG (10 µM) but not in untreated cells. **(B)** Representative fluorescence image showing time course of RACK1 condensate formation in HCT-8 cells expressing RACK1-GFP stimulated with IL-1β (10 ng/μL). **(C)** Fluorescence recovery after after photobleaching (FRAP) time course of RACK1-GFP condensates after IL-1β (10 ng/μL) stimulation(orange) and unstimulated cytosolic RACK1-GFP (black). The solid line represents the mean and the dash line represents the error bounds defined by the standard deviation. **(D)** Diffusion properties of RACK1 puncta upon stimulation with IL-1β (10 ng/μL) relative to control. (n ≥3) **(E)** Representative fluorescence image showing colocalization of RACK1-GFP and INAVA-mcherry in HCT-8 cells upon stimulation with IL-1β (10 ng/μL) or with 17-AAG (10 µM). **(F)** Pearson correlation of coefficient to quantify colocalization between RACK1-GFP and INAVA-mcherry in HCT-8 cells upon stimulation with IL-1β (10 ng/μL) or with 17-AAG (10 µM). (n=2) **(G)** Representative co-immunoprecipitation blots showing interaction between RACK1 and GFP at two different timepoints in unstimulated or when stimulated with IL-1β (10 ng/μL) or with 17-AAG (10 µM) **(H)** Quantification of (G). (n=2) **(I)** Time course of change in cytosolic reactive oxygen species measured by normalized fluorescence change of CM-H_2_-DCFDA in HCT8 cells after IL-1β (10 ng/μL) or with 17-AAG (10 µM) (n=2) **(J)** Representative fluorescence image of Caco-2 cells, untreated or stimulated with IL-1β (10 ng/μL) and/or Nox1 inhibitor for 1 hour and stained with DCP-Rho1(10DμM). **(K)** Representative ratiometric fluorescence image of HCT-8 cells overexpressing RACK1-HyPer3. The intensity is the ratio of fluorescence signals excited at 488 nm and 405 nm. **(L)** Quantification of intensity ratio described in (K) as a function of time. (n ≥3) *p < 0.05, **p < 0.01, ***p < 0.001

To assess whether RACK1 puncta are phase-separated structures (21), we conducted fluorescence recovery after photobleaching (FRAP) experiments to measure diffusion properties (Figure 3C). We observed that compared to control, these stress-induced puncta have a lower molecular diffusion coefficient and mobile fraction (Figure 3D, S2B). Following 17-AAG stimulation, puncta displayed reduced diffusion coefficients over time suggesting aggregate formation at longer times (Figure S2C). Analysis of the RACK1 secondary structure indicated no major intrinsically disordered regions suggesting a low propensity to induce condensate formation (Figure S2C). We hypothesized that RACK1 could exist in condensates as a client of scaffolding condensate forming proteins.(22) In a previous work, we had observed that Innate Immunity Activator (INAVA), a highly disordered protein (Figure S2D-F), forms condensates upon exposure to IL-1β in epithelial cells.(23) To test whether INAVA could be a potential condensate forming protein that recruits RACK1 in epithelial cells we localized both INAVA and RACK1. Under both IL-1β and 17-AAG treatment, RACK1 and INAVA were found to co-localize in puncta (Figure 3E, 3F). Co-immunoprecipitation studies indicated that interactions between RACK1 and INAVA progressed with time following stimulation (Figure 3G). We observed that with both IL-1β and 17-AAG treatment, RACK1 was recruited to INAVA, peaking at around 30 minutes, and temporally mirroring the dynamics of cellular ROS (Figure 3H, 3I). In earlier work, RACK1 was shown to localize to stress granules following cellular insult (24). Colocalization experiments of RACK1 with the stress granule marker, G3BP showed large cell to cell variation under different stressors (Fig S2G, H). This variation in colocalization of the two proteins by treatment was seen with 17-AAG inducing relatively high co-localization in contrast to IL-1β which showed limited colocalization.

We then assessed for global localization of redox-modified proteins using the fluorescent indicator DCP-Rho which marks oxidized cysteine sulfenic-acid containing sites in proteins. Here upon IL-1β stimulation we observed the appearance of distinct puncta distributed throughout the cell (Figure 3J), suggesting the possibility that these puncta may be phase separated condensates that are enriched for protein oxidation. ROS and protein redox status are known to be very tightly controlled within cells, with the general cytosolic environment thought to be highly reducing to avoid non-specific oxidation.(25) Given our finding of redox regulation of RACK1 and the temporal dynamics of ROS and puncta formation, we assessed whether RACK1 containing condensates may be sites of spatially restricted elevated H_2_O_2_. Using RACK1 tagged with a genetically encoded ratiometric H_2_O_2_ sensor (HyPer3) we observed that RACK1 containing condensates exhibited a time dependent increase in relative oxidation compared to the cytosol providing evidence for condensates as a cellular redox-privileged site (Figure 3G).

### RACK1 puncta formation and response to redox are evolutionarily conserved

Given that RACK1 signaling and its biophysical response to oxidation appear under seemingly diverse stimuli, we speculated that cysteine residues within core WD repeat domains may be a highly conserved feature. Six of the eight cysteines in human RACK1 are in the WD repeats and are highly evolutionarily conserved with C286 among the most conserved (Figure 4A). The only model eukaryote where C286 was not conserved was in *Saccharomyces cerevisiae* although, *Saccharomyces pombe* is similar to higher eukaryotes possessing the cysteine at the 286 position.

**Figure 4.**
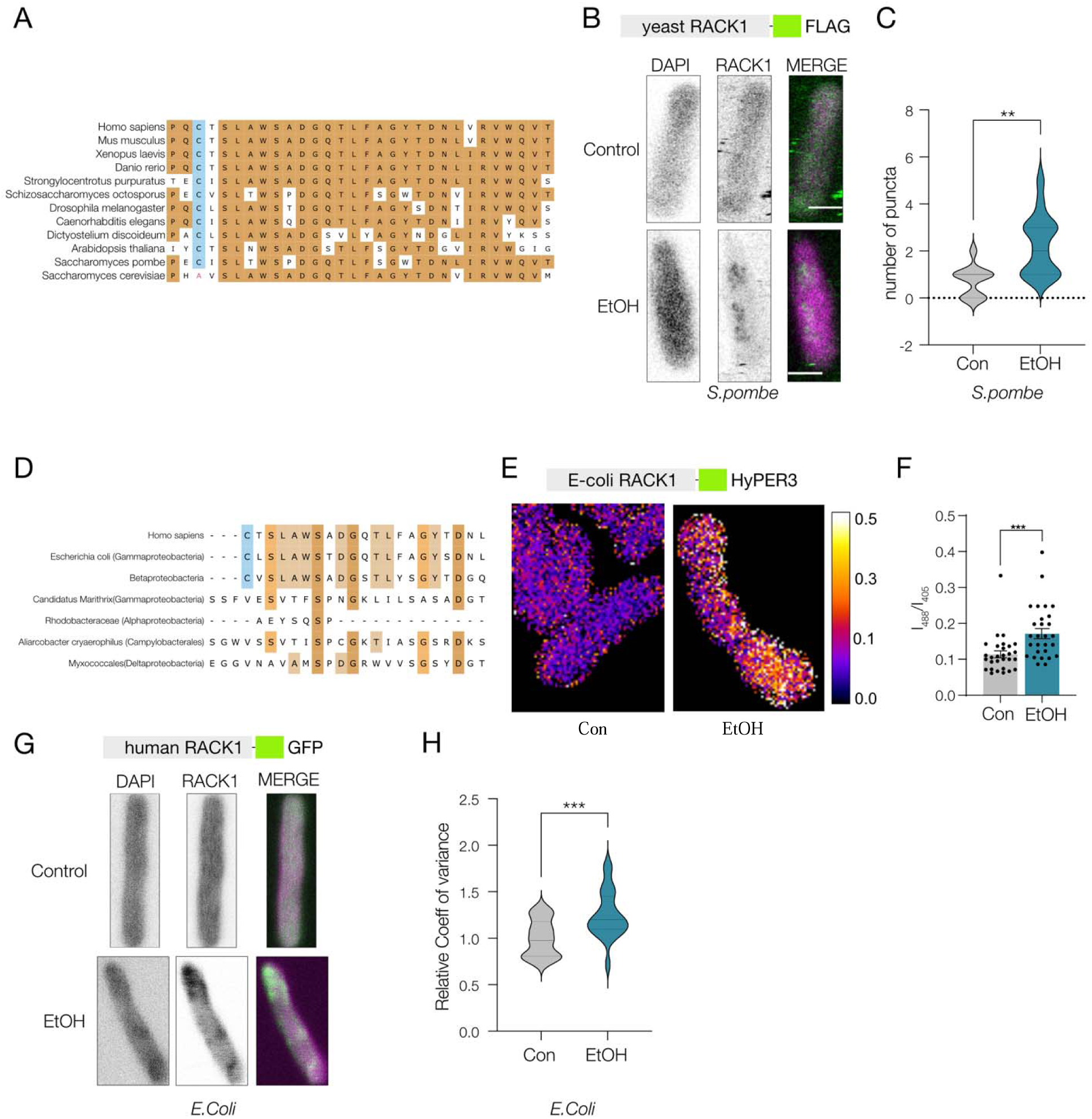
RACK1 puncta formation and response to redox are evolutionarily conserved. **(A)** Multiple Sequence alignment of amino acids proximal to cysteine 286 in eukaryotes. **(B)** Representative immunofluorescence image of fission yeast cells expressing Flag-cpc2 and treated with either 10% ethanol or media control for 10 minutes. **(C)** Quantification of number of puncta per cell in control and ethanol treated yeast. (n=3) **(D)** Multiple sequence alignment of the amino acids proximal to cysteine 286 in human RACK1 against bacterial homologs. **(E)** Representative ratiometric fluorescence image of E. *coli* expressing bacterial analog of RACK1 tagged to Hyper3. **(F)** Quantification of intensity ratio of E. *coli* expressing bacterial analog of RACK1 tagged to Hyper3 when treated with either 10% ethanol or media control. (n=3) **(G)** Representative fluorescence image of E. *coli* expressing human RACK1 protein tagged with GFP treated with either 10% ethanol or media control for 10 minutes. **(H)** Quantification of normalized coefficient of variance in GFP signal as a measure of heterogeneity of RACK1 signal in the cytosol. (n=3) *p < 0.05, **p < 0.01, ***p < 0.001

Given the high sequence similarity observed between evolutionarily distant species (Figure S3A), we hypothesized that RACK1 has conserved functional responses in lower eukaryotes and tested if the RACK1 homolog in *Saccharomyces pombe*, CPC2 also localize to puncta under stress. Fission yeast (h-leu1-32 ura4-D18 Flag-cpc2) were stressed with an ethanol shock and assessed for CPC2 localization. Under stress CPC2 also partitioned into punctate structures, with a significantly increased number of CPC2 puncta than controls, indicating that in lower eukaryotes, RACK1 also forms stress-induced condensate-like structures (Figure 4B, 4C).

Since all eukaryotes have the RACK1 gene, we investigated whether protein orthologs of RACK1 may be present in either prokaryotes or archaea and have functionally analogous stress responses. Interrogation of sequence similarity showed that while RACK1 orthologs were absent in archaea, we found that several bacterial species had proteins with high similarity to human RACK1 (Figure S3B, C). Of all bacterial species, cyanobacteria have the most abundant and diverse set of RACK1-like proteins (Figure S3C, S3E), but do not contain a conserved cysteine analogous to C286 in the WD7 repeat domain. However, several species of proteobacteria, including *E. coli* and Bacillota species, possesses WD-40 like proteins similar to RACK1 and contain a conserved cysteine in a position analogous to C286 in hRACK1 (Figure 4D, S3D).

To assess analogous function in prokaryotes we transformed *E. coli* strain BL21 with a HyPer3 tagged bacterial analog of RACK1 and subjected to ethanol shock to induce environmental stress. Imaging of *E. coli* subjected to ethanol indicated localized areas of increased H_2_O_2_ relative to control, suggesting that similar to human and yeast cells, bacteria respond to environmental stress with localized increased oxidation in the vicinity of RACK1 (Figure 4E, F). We also expressed human GFP-RACK1 in *E. coli* and assessed protein localization under ethanol stress. We again observed that RACK1 are localized to heterogenous puncta (Figure 4G, 4H). Our results suggest that similar to eukaryotic cells RACK1 can associate with redox-enriched condensate-like structures in bacteria when subjected to stress.

### Oxidation of RACK1 in condensates alters protein translation

To investigate the consequence of stress induced RACK1 oxidation and sequestration into condensates we interrogated cellular modules altered following RACK1 cysteine oxidation. We conducted pathway analysis of proteins identified in our redox proteomics screen and found the Gene Ontology terms corresponding to proteosome and translation to be significantly enriched (Figure 5A). Regulation of protein translation is a major and conserved cellular adaptation to environmental stress.(26) Since RACK1 is a known protein component of the 40S subunit of ribosomes we investigated whether redox regulation of RACK1 may be involved in stress induced changes in protein translation. To test this, we conducted puromycin incorporation experiments to determine if loss of cysteine oxidation alters cellular translational responses. We observed that following 17-AAG induced stress, cells overexpressing WT RACK1 exhibited significant translation inhibition as demonstrated by reduced puromycin incorporation relative to untreated controls (Figure 5B, 5C). In contrast, cells containing the RACK1-C-all mutant had significantly lower translation inhibition relative to wildtype following 17-AAG induced stress.

**Figure 5.**
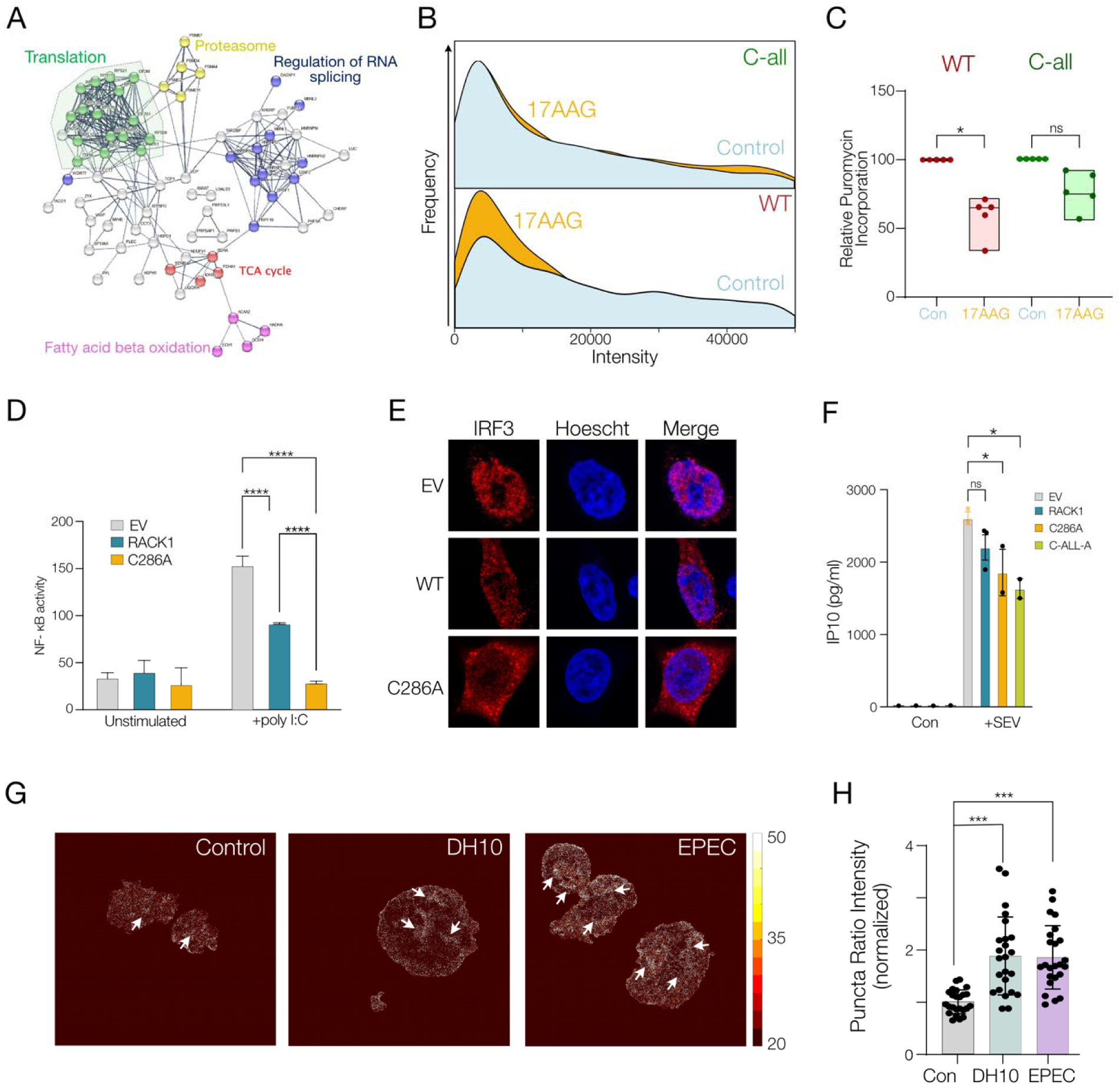
Oxidation of RACK1 in condensates is critical for cellular function. (**A**) String network analysis of redox-proteomics screen colored by manually curated subset of significant functional modules. (**B**) Representative flow cytometry data showing normalized intensity of puromycin incorporation in HCT-8 cells expressing either Rack-1 WT mCherry (top) or Rack-1 Call mCherry (bottom) treated with 17-AAG (10 µM, 10 minutes, orange curve) or untreated (blue). **(C)** Puromycin incorporation in HCT-8 cells expressing either Rack-1 WT mCherry or Rack-1 C-all mCherry treated with 17-AAG (10 µM, 10 minutes) normalized to untreated. (n=5). **(D)** NF-κB activity in HEK cells overexpressing empty vector, RACK1-WT or RACK1-C all stimulated with LPS (10 mg/ml). (n≥3) **(E)** Representative fluorescence image showing IRF3 translocation in HEK cells overexpressing empty vector, RACK1-WT or RACK1-C-all stimulated with poly I:C (10µg/ml) **(F)** IP10 production following infection of HEK293T cells overexpressing empty vector, RACK1-WT, RACK1-C286A or RACK1-C-all mutant with Sendai virus (MOI = 1, 1h) (n≥2)**(G)** Representative ratiometric fluorescence image of human duodenal cells transfected with Rack1-Hyper3 and exposed to E. *coli* strains—pathogenic EPEC O127:H6, DB10, or uninfected. **(H)** Quantification of number of Mean Puncta Intensity ratio (405/488) per transfected cells. (n=3) *p < 0.05, **p < 0.01, ***p < 0.001

### RACK1 oxidation and condensate formation in response to pathogen induced stress

Since redox-regulation of RACK1 appeared to be an evolutionarily conserved response to stress we investigated whether RACK1 function and oxidation may be a generalized stress-effector system in response to pathogens in human cells. We determined the oxidation percentage of cysteines by performing OxICAT analysis of wildtype and NOX1^-/-^ colonoids following stimulation of the synthetic dsRNA ligand and viral mimetic poly I:C.(16, 27, 28) OxICAT analysis identified cysteine oxidation status in unstimulated and poly I:C stimulated WT and NOX1^-/-^ colonoids (Figure S4A, B). Identified cysteines in RACK1 were found to be amongst the most highly differentially oxidized following treatment with poly I:C. Functionally, poly I:C induced NFκB activation was found to be inhibited by RACK1 and further depressed by RACK1-C286A (Figure 5D). Consistent with a role in both viral and bacterial signaling, IRF3 translocation into the nucleus in response to poly I:C was also lowered in cells expressing RACK1-C286A (Figure 5E). Infection of cells with Sendai Virus induces a robust interferon-stimulated response in host cells as shown by secretion of the chemokine IP10. Cells expressing RACK1 cysteine mutants, C286A or C-All showed significantly lower IP10 secretion (Figure 5F) suggesting redox-regulation of RACK1 is a generalized pathogen stress response. Finally, to assess RACK1 oxidation for pathogen responses in primary human cells, we treated patient-derived duodenal cells grown as monolayer patches expressing Hyper3-RACK1 with either *E. coli* DH10B or Enteropathogenic *E. coli* (EPEC). Here we observed that both DH10B and EPEC infection of intestinal epithelial cells induced increased RACK1 puncta formation relative to control media treated cells. RACK1 puncta but not cytosolic RACK1 in *E. coli* infected cells had significantly increased H_2_O_2_ relative to uninfected cells (Figure 5G, H, S4C, S4D).

## Discussion

In this manuscript, we report an ancient stress sensor-effector system comprising of a highly conserved protein, RACK1 as a key transducer of cellular redox responses. We show that RACK1 populates membrane-less condensates which form a spatiotemporal platform to enable stabilization of cysteine oxidation and allow sensing of short-lived redox signals within the reducing cytosolic environment of the cell.

Initial conditions on Earth were highly anoxic with oxygen being toxic to organisms. Slowly, cells adapted to consume oxygen although the cytoplasm itself remained reductive. Reactive oxygen species such as H_2_O_2_ with a short half-life in the cell evolved as signaling molecules to regulate complex cellular processes through post-translational modifications of amino acids, most prominently cysteines.(29) Numerous examples of the influence of redox signaling have emerged across kingdoms of life, ranging from abiotic stress in plants to immune responses in humans.(30, 31) Given the potential for cellular damage via H_2_O_2_, cells maintain a generally highly reductive cytosolic environment to prevent non-specific protein oxidation. Targeted oxidative modification of cysteine residues and thiol switch formation to enable signaling is dependent on the local environment around proteins. Environmental conditions alter the chemical state stability of thiol switches, resulting in a wide range of reactivity towards H_2_O_2_, making local H_2_O_2_ concentrations a key and protein-specific determinant of oxidative modification.(32) Therefore, the enabling of oxidative protein modifications requires the generation of spatially sequestered environments that allow for localized pools of H_2_O_2_ that are shielded from the action of cytosolic reducing equivalents. In this context our finding that RACK1 condensates exhibit a spatially compartmentalized redox environment provides evidence for a redox signaling regulatory mechanism with multiple implications for the control of cellular functions.

Our finding that oxidation of RACK1 is important for cell signaling stemmed from our proteomic screen and was validated by cysteine mutation studies showing modulation of NFκB activity as well as responses to diverse environmental stress signals, suggesting a generalized effector pathway. RACK1 is a multifunction cellular signaling protein that has previously been associated with stress response in cells,(24, 33) often through interactions with stress granules.(34, 35) This recruitment into stress granules and function is associated with interaction with binding to ribosomes and through post-translational modification including MARylation.(33) In keeping with these findings, we found that a variety of stressors induced RACK1 puncta formation in the cytosol of mammalian cells. We show that these phase-separated membrane-less condensates form a localized redox-privileged niche environment. Importantly, stress-induced RACK1-enriched puncta were observed in yeast and bacteria suggesting an ancient origin of this mechanism consistent with the early role of condensates as functional entities given that amino acid-RNA coacervates are observed across multiple domains of life.(36) Cell-free *in vitro* studies along with modeling suggest that the interfacial properties of condensates alter internal electric potential which in turn promote redox reactions.(37) Studies have shown that in different cellular systems, the functional state of condensate-associated proteins can be modulated by the redox state of amino acids, in the context of phase separated granules.(38–40) In our studies, we show that RACK1 resides in a relatively oxidizing niche environment within condensates, that likely maintains and facilitates functions dependent on its cysteine modifications. RACK1 has a well appreciated function as part of the 40S ribosome subunit and our redox proteomic analysis indicated a distinct signature related to translation regulation.(41) We show that RACK1 appears to connect stress induction to translation inhibition via redox-sensitive modifications within the condensate environment consistent with a major known function of stress granules.(40, 42)

We postulate that signaling specificity of this redox stress-effector system is likely generated by the association of the protein sensor in the system - RACK1 - with different protein and RNA partners. In this setting RACK1 by itself is likely not a driver of condensate formation but exists as a client to other primary condensate forming proteins in cells. RACK1 is a generally ubiquitously expressed protein, and we speculate that there are cell and context specific proteins that associate and drive RACK1 condensate formation to enable a wide diversity of signal propagation. In intestinal epithelial cells, this protein partner appears to be INAVA, an intrinsically disordered protein, that has functions in both barrier function and innate immunity, although we cannot ascertain if the interaction is initiated in the condensate itself or occurs in the cytosol and then trafficked to the condensate.(23) We also hypothesize that there could be other more ubiquitously expressed proteins, including G3BP that functionally co-localize within these condensates depending on context and type of stress. Other potential candidates that may drive RACK1 condensate formation include SRSF1 and SFPQ proteins. Concurrent studies using methionine substitution in relation to H_2_O_2_ responses, showed these two proteins are redox-responsive and that stress granule responses involved in amyotrophic lateral sclerosis may be dependent on redox state.(39)

RACK1 appears to be a conserved node within redox signaling networks with particular preservation of the key cysteine C286 which is unique to WD7 (Figure S3B). Overall, the similarity of the RACK1 ortholog CPC2 protein in *Sacchromyces pombe* is almost 80% compared to human RACK1. This makes RACK1 more conserved than other prototypically highly conserved proteins such as small ribosomal subunit protein, 40S ribosomal protein SA (RPS-A), which has a sequence similarity of 70% between *pombe* and human. In bacteria we found several species of proteobacteria, including *E*. *coli* and Bacillota, possess WD40-like proteins similar to RACK1 with a conserved cysteine in a position analogous to C286. Interestingly, the putative RACK1-like protein in *E*. *coli* has over 70% sequence identity with human RACK1, although not all proteobacteria analyzed have this conserved cysteine (Figure S3D). Interestingly, of all bacterial species, cyanobacteria have the most abundant and variable RACK1-like proteins (Figure S3E). Despite a careful search for cysteines in a WD7 repeat domain, we were unable to find any example of this in cyanobacteria. Given the sequence similarity between higher proteobacteria and eukaryotes, we speculate that eukaryotic RACK1 had a bacterial origin arising potentially through an early horizontal gene transfer event similar to mitochondria. However, we hypothesize that this occurred distinctly from mitochondrial gene transfer since amitochondrial eukaryotes such as Entamoeba have a RACK1-like protein with a C286 analogue (Figure S3F), but do not have mitochondrial genes such as GSH.

In summary, we find that RACK1 cysteine oxidation within redox-privileged condensates is an ancient and relatively ubiquitous sensor-effector system that operates to transduce environmental information to cellular stress responses. We show that this system operates for a variety of innate immune responses in mucosal epithelial cells and therefore may be important for pathological responses in a wide range of disease states.

## Acknowledgments

We would like to thank the Kagan and Garber lab for sharing reagents used in this paper. We would like to thank Ajit Joglekar for help with the yeast experiment.

## Funding

National Institutes of Health grant R35GM142683 (JRT), R35GM134964 (EW)

### Author contributions

Conceptualization: EN, JRT

Methodology: EN, SS, KR, WIL, EW, JRT

Investigation: EN, XC, SS, AM, VS, QP, IM, KR, JRT

Funding acquisition: JRT, WIL, EW

Project administration: JRT

Supervision: EW, JRT

Writing – original draft: EN, KR, JRT

Writing – review & editing: EN, SS, XC, KR, WIL, EW, JRT

## Data and materials availability

All data are available in the main text or the supplementary materials.

## Online Methods

### Cell culture

All cells were maintained at 37°C in a humidified incubator with 5% CO. Caco-2 bbe, HEK293T, and HCT8 cells were cultured in Dulbecco’s Modified Eagle Medium (DMEM) supplemented with 10% fetal bovine serum (FBS) and 1× Antibiotic-Antimycotic (Gibco). Mouse intestinal enteroids and human colonic organoids were maintained in mouse and human organoid medium. For redox experiments, organoids were cultured in antioxidant-free medium for at least one passage prior to use. Human and mouse enteroids were mechanically dissociated and treated with cell recovery solution (Corning) and embedded in Matrigel. Sendai virus was shared by the Kagan lab. EPEC strains were donated by the Garber lab

### Organoid isolation

Mouse colonic crypts were isolated from age-matched wild-type and Nox1 / mice. Animals were euthanized by CO inhalation and colons excised, opened longitudinally, and washed thoroughly in ice-cold PBS. Tissues were incubated in 10 mM EDTA in PBS at 4°C with agitation for 45 min to release crypts, followed by filtration through a 40-μm strainer and centrifugation. Crypts were washed and embedded in Matrigel (Corning), plated in 24-well plates, and overlaid with mouse organoid medium. Organoids were passaged bi-weekly.

### qPCR

Total RNA of cells was extracted with the RNeasy Mini Kit (QIAGEN). CDNA was prepared from total RNA with the SuperScript First-Strand Synthesis System (Invitrogen) with the oligo (dT)12–18 primers, according to the manufacturer’s instructions. RNA samples were treated with DNase I (Invitrogen) for the elimination of genomic DNA. Cytokine mRNA expression was determined by real-time qPCR with relative quantitation by the comparative threshold cycle number (Ct) method, using an iCycler and SYBR Green Ready-Mix (Bio-Rad) and normalized by a housekeeping gene (β-actin or GAPDH).

### Strain construction

Full-length human RACK1 (WT), C249A, C286A, C249A/C286A (double mutant), and all-cysteine-to-alanine (Call) constructs were synthesized (IDT) with C-terminal HA tags and cloned into pcDNA4 using HindIII and NotI. For immunofluorescence experiments mKate-tagged versions (WT, C286A, Call) were cloned into pLVX using XmaI and XbaI. E. *coli* Rack1 constructs were synthesized and cloned into pET28 using SacI and NotI. The pEGFP-N1-Rack1 plasmid was obtained from Addgene (#41088).

### Gene expression quantification

Total RNA was extracted using the RNeasy Mini Kit (Qiagen) and reverse transcribed with the QuantiTect Reverse Transcription Kit (Qiagen). Gene expression was quantified via qPCR using SsoAdvanced SYBR Green Supermix (Bio-Rad) with primers from IDT. Target expression was normalized to β-actin and quantified as the 2^–ΔΔCt.

### Rack I knockdown by siRNA

For siRNA knockdown, HEK293 cells were transfected with pooled RACK1-targeting siRNAs (Qiagen, Cat. No. 1027416) or control siRNAs (AllStars Negative and Cell Death Control). Lipofectamine RNAiMAX (Thermo Fisher) was used for transfection. Cells were harvested after 48 or 72 hours, lysed in RIPA buffer, and protein concentrations measured by BCA assay. Western blots were probed with anti-RACK1 and anti–β-actin antibodies and visualized by chemiluminescence.

### Co-Immunoprecipitation for analysis of protein interactions

Immunoprecipitations were performed in HEK293T cells treated with IL-1β (10 ng/μL, 1 hr.) or control using methods modified from Tan and Kagan (2018).(43) Cells were lysed in RIPA buffer containing protease and phosphatase inhibitors. Three quarters of the lysates (centrifugation at 16,000 × g for 15 minutes at 4 °C) were incubated with 1ug of primary antibodies and Protein G Sepharose overnight at 4 °C, and the rest of 4× Laemmli loading buffer, boiled at 100 °C for 5 minutes and stored at −20 °C to serve as input. Complexes were washed extensively, eluted in Laemmli buffer, and analyzed for interacting proteins by SDS–PAGE and immunoblotting.

### Biotin Switch for analysis of oxidized proteins

Biotin-switch assays were adapted from Yang et al. (2018). (44) Caco-2 cells stimulated with IL-1β were lysed and biotin alkylated in NEM-containing RIPA buffer, clarified, and desalted. Protein amounts were normalized using BCA assay and, sodium arsenite (200mM) and biotin-maleimide (1mM) was added to selectively reduce Sulfenic acid groups and label newly synthesized thiols. Labeled proteins were enriched using NeutrAvidin beads, eluted, and analyzed by SDS–PAGE and Western blot.

### Sendai virus (SeV) Infection of HEK

For Sendai virus (SeV) infection, HEK293T cells overexpressing empty vector or RACK1-Call mutant were infected with SeV (MOI = 1; sourced from Canteli Allantoic Fluid Lot 7Y111007, Charles River Labs) for 1 hour in DMEM + 1% FBS, washed, and incubated for 6 h to overnight. The supernatant from the infected HEK Blue cells was used to measure the IP-10 production with Human CXCL10/IP-10 Immunoassay (R&D System, DIP100) following the manufacturer’s instructions. The optical density of each well was measured using Tecan Spark 10M Multi-Mode Plate Reader set to 450 nm and 570 nm. The optical density at 570 nm was subtracted from measurement at 450 nm to obtain normalized values. All infection steps were performed in a biosafety level 2 (BSL-2) containment environment with appropriate personal protective equipment (PPE).

### siRNA Transfection and Immunoblotting

HEK293 cells were transfected at ∼80% confluency with pooled RACK1-targeting siRNAs (Qiagen, Cat. No. 1027416) or control siRNAs; AllStars Hs Cell Death Control and AllStars Negative Control siRNAs (Qiagen) using Lipofectamine RNAiMAX (Thermo Fisher) in Opti-MEM (Gibco). Transfections were performed in 6-well plates and incubated at 37 °C for 48 or 72 h. Cells were lysed in RIPA buffer, and protein concentration was measured by BCA assay. Equal protein amounts were resolved by SDS–PAGE, transferred to PVDF membranes, and probed with anti-RACK1 and anti–β-actin antibodies for chemiluminescent detection.

### Evaluation of protein oxidation in Caco-2Bbe Cells Using DCP-Rho

Caco-2 cells were grown to ∼70–80% confluency in 24-well plates. To assess protein sulfenylation, cells were treated with or without interleukin-1β (10 ng/mL) for 1 hour. DCP-Rho1 (final concentration 10 μM; Kerafast) was added directly to the culture medium during the final 10 minutes of the IL-1β stimulation period. Following probe incubation, cells were washed three times with ice-cold phosphate-buffered saline (PBS) to remove unbound probe and then fixed for imaging. All treatments were performed in triplicate. Fluorescence was detected by confocal microscopy using excitation/emission settings of 552/575 nm.

### Transfection and Bacterial Infection of Human Duodenal Organoid Monolayers

Human duodenal organoids (WT) were cultured to stem-like phase in Matrigel domes (12 wells total). For transfection, Matrigel was removed, and organoids were dissociated into fragments with TrypLE Express (2 mL, 10 minutes at 37°C). Dissociation was quenched with 10% FBS-containing medium and suspension was filtered through a 40 μm strainer. Cells were washed and approximately 1 × 10 to 5 × 10 cells were resuspended in Opti-MEM (100 μL). 10 μg Rack1-Hyper3 or empty vector plasmid DNA (DNA volume ≤10% of total) was added to cells and the mix was transferred to an electroporation cuvette and electroporated using the (NEPA21, Bulldog) with optimized settings based on Manufacturers protocol. The poring pulse phase and the transfer pulse settings were as follows: Voltage (175 and 20V), Pulse Length (5 and 50 msec) and Number of Pulses (2 and 5). The final impedance was verified as to be between 30-55 Ω. Following recovery in Opti-MEM, cells were seeded onto Matrigel coated glass-bottom plates and cultured for one to two weeks to form monolayers.

For infection, monolayers were exposed to *E. coli* strains—pathogenic EPEC O127:H6 (JPN15) or control DH10B. Infections were carried out with 5 × 10 bacteria per well for 6 hours to overnight. (45) Ratiometric redox imaging was then conducted. Confocal images were acquired using Zeiss LSM 880, a 63X objective (NA 1.4). Two channels were imaged (1024X1024 pixels at 43.9 nm resolution). For ratiometric calculation of Hyper-3, two excitation lasers (405 nm and 488 nm) were used to sequentially image an emission channel (500-530 nm). Images were analyzed using custom written MATLAB code to generate the redox ratio metric images (I405/I488). Each channel was first eroded using a structural element of 10 pixels. To determine the cell boundaries, a composite image of both the channels were added, gaussian- and diffuse filtered, and then multi-otsu thresholded. The binary image was filtered for small particles and the final binary image used to determine the mask for both the channels. The ratio of the channels was calculated and any ratio greater than 20 was set to 0 to prevent low signal regions.

### Luciferase reporter assay

15000 viable HEK 293 or HEK Blue cells/100µl DMEM/FBS were seeded in each well of a 96-well plate and grown at 37°C overnight. Cells were then transfected using transfection reagent lipofectamine 3000 (Thermo Fisher, L3000001) following the manufacturer’s instructions. Typically, we transfected 200 ng DNA per well of a 96-well plate, including 35ng pNL3.2.NF-kB-RE [Nluc/NF-kB-RE/Hygro] Vector (Promega, N1111) and 165 ng pcDNA4.0-empty vector, RACK1-WT, RACK1-C286A or RACK1-C-ALL plasmid. 24 hours after transfection, medium was changed and cells were pretreated w/wo 5mM NAC or 1mM Rotenone, then HEK 293 cells were stimulated with IL-1β, TNFα or LPS, HEK Blues cells were stimulated with poly I:C in indicated doses for 6 hours. Nano-Glo® Luciferase Assay System (Promega, N1130) was used to detect luciferase activity in a lytic method. Briefly, we prepared the desired amount of reconstituted Nano-Glo® Luciferase Assay Reagent by combining one volume of Nano-Glo® Luciferase Assay Substrate with 50 volumes of Nano-Glo® Luciferase Assay Buffer. All assay components (reagent and sample) were allowed to equilibrate to room temperature prior to measurement. For 96-well plates, typically 100μl of combined reagent was added to the cells grown in 100μl of medium. Luminescence was measured by Tecan Spark 10M Multi-Mode Plate Reader after an incubation time of at least 3 minutes.

### Transfection, Immunofluorescence staining and microscopy

For p65 and IRF3 translocation microscopy, we seeded HEK 293 or HEK Blue cells in a 24-well glass-bottom plate and placed the cells in a tissue culture incubator at 37°C overnight to get 70% confluency the next day. We then transfected the cells using Lipofectamine 3000 (Thermo Fisher, L3000001) following the manufacturer’s instructions. Typically, we transfected 1000 ng DNA of pcDNA4.0-empty vector, RACK1-WT or RACK1-C286A per well of a 24-well plate. 24 hours after transfection, medium was changed and HEK 293 cells were stimulated with 10 ng/ml IL-1β for 30min, HEK Blue cells were stimulated with 10 µg/ml poly I:C for 2h separately. The cells were then fixed with 4% paraformaldehyde diluted in PBS for 15min at room temperature and washed with PBS three times. The samples were blocked with 10% donkey serum in PBS for 1 h at 4 °C in a humidified chamber, incubated with the indicated primary antibodies diluted in blocking buffer at 4 °C overnight, washed twice and then stained with fluorescent secondary antibodies diluted in blocking buffer for 2 h at room temperature. Finally, cells were washed three times and incubated with PBS containing Hoechst at a 1:1,000 dilution for 10min and then mounted with ProLong Dimond Antifade Mountant (Thermo Fisher Scientific). Images were taken from one or more sections with a 20× objective on a Zeiss 880 inverted confocal microscope. The following primary antibodies and dilutions were used: p65 (CST, 8242S, 1:1000) and IRF3 (CST, 11904S, 1:1000). For the colocalization experiments with G3BP, confocal images were acquired using Abberior Stedycon system a 100X objective (NA 1.4). Three channels, corresponding to DAPI, G3BP (Secondary Antibody 488) and RACK1-mcherry were imaged at 50nm resolution. Pearson Correlation Coefficient (ICY, bioimage analysis platform) was measured for each cell, whose regions were manually annotated. Three independent experiments were conducted for both Il1β and 17-AAG.

### Bioinformatics Analysis

RACK1 sequence analysis was performed using either MolEvolvR, using custom R programs utilizing Biostrings, spiralize, MSA and ggMSA packages and Unipro UGENE (ver 44.0). (46–48) Blast Analysis was conducted using the Uniprot website.(49) MSA package was used to generate conservation scores. The structure prediction was conducted using Alphafold.((50) Disorder prediction score of the different proteins was calculated using IUPred 3 webtool.(51, 52)

### Yeast Puncta Experiments

h-leu1-32 ura4-D18 Flag-cpc2 yeast strains were obtained from Yeast Genetic Resource Center (YGRC), Osaka, Japan. Yeast strains were grown in EMM media with no nitrogen (no NH_4_CL, glutamate) and supplemented with L-Glutamic acid (20 mM), Leucine (200mg/L) and uridine (200mg/L). Cells were grown overnight and treated with either 10% ethanol or media control for 10 minutes, fixed with 4% PFA and incubated in Blocking Buffer containing 0.1% Triton X-100 and 2% BSA. The cells were then incubated with Anti-Flag Primary Antibody (F7425 Sigma), washed with blocking buffer and incubated with Secondary Antibody against the FLAG antibody, along with Hoech st. Cells were imaged using Abberior Stedycon system a 100X objective (NA 1.4) and the puncta was manually counted by a researcher who was blinded.

### Pulldown Assays

HCT8 cells expressing doxycycline (dox)-inducible myc-ARNO and HA-INAVA tags were seeded onto 10 cm culture dishes and incubated overnight to allow the cells to reach approximately 70–80% confluence by the following day. At the time of plating, 1 µg/mL of doxycycline was added to designated control dishes to induce expression. The next day, cells were treated with one of the following conditions for various time intervals (10, 30, 60, or 120 minutes): Full growth media (DMEM supplemented with 10% FBS and penicillin/streptomycin); IL-1β at a final concentration of 10 ng/mL; or 17-AAG at 10 µM. Following treatment, cells were washed three times with cold PBS and lysed in RIPA buffer for 30 minutes at 4°C. The lysates were then centrifuged at 5,000 × g for 15 minutes at 4°C, after which the resulting supernatants were collected and split into two portions for use as input and co-immunoprecipitation (Co-IP) samples. Protein G Sepharose bead slurry (pre-washed in PBS and resuspended in RIPA buffer) was incubated with 1 µg of anti-HA antibody and added to the Co-IP fractions. Samples were rotated overnight at 4°C on a nutator. Beads were then pelleted by centrifugation at 3,000 × g for 30 seconds at 4°C, washed three times with cold PBS, and resuspended in Laemmli buffer. Both input and Co-IP samples were then boiled at 95°C for 5 minutes prior to western blot analysis.

### FACS-Based Reactive Oxygen Species (ROS) Assay

Wild-type HCT8 cells were plated in 24-well plates two days before the assay to ensure they reached 70–80% confluence at the time of treatment. On the day of the assay, cells were washed twice with PBS and subsequently incubated with 5 µM CM-H DCFDA, a fluorescent ROS indicator, for 10 minutes at 4°C. Following labeling, the cells were washed twice with complete growth media and then exposed to one of the following conditions for 0, 10, 30, 60, or 120 minutes: Full growth media (DMEM with 10% FBS and pen/strep); IL-1β at 10 ng/mL; or 17-AAG at 10 µM. After treatment, cells were washed twice with PBS, trypsinized, and filtered through a FACS tube strainer to ensure a single-cell suspension. Samples were then analyzed for ROS levels using a BD FACSCanto II flow cytometer.

### Colocalization Imaging of INAVA and RACK1

HCT8 cells stably overexpressing INAVA-mCherry were transiently transfected with a RACK1-GFP plasmid using Lipofectamine 3000 in 24-well glass-bottom plates. Transfections were performed overnight according to the manufacturer’s protocol. The next day, cells were treated for 30 minutes with one of the following: Full growth media; IL-1β (10 ng/mL), or 17-AAG (10 µM). After treatment, cells were fixed using 4% paraformaldehyde (PFA) for 15 minutes at room temperature. Confocal microscopy was then used to image the samples and assess colocalization between INAVA and RACK1.

### Puromycin incorporation assay

HCT-8 cells expressing either Rack-1 WT mCherry or Rack-1 Call mCherry were washed twice in full media and treated with either 17-AAG at 10 µM or untreated for 10 min. Cells were subsequently washed in full media and then supplemented with puromycin [10ugml] for 10 min. Puromycin was washed off and cells were incubated further in 17-AAG, IL-1, cycloheximide or left untreated for another 50 min.

Cells were then washed with PBS and cells were dissociated using cell dissociation buffer (Thermo, Catalog number 13151014). Cells were fixed and permeabilized using cell Fix and Perm medium (Invitrogen, Catalog number GAS002S100) according to manufacturer’s instruction. Cells were then stained with Alexa647 conjugated anti Puromycin antibody [1:2500] for 30 min on ice and washed 3x with PBS. Cells were analyzed using flow cytometry.

### Fluorescence Recovery After Photobleaching

FRAP experiments were performed on a Zeiss 880 laser scanning confocal microscope using the in-built FRAP module within the microscope control software Zen Black. HCT8 expressing GFP-tagged RACK1were plated on a 35 mm μ-Dish (Ibidi; Munich, Germany). As indicated, cells were cotreated with IL-1β (10 ng/mL), or 17-AAG (10 µM). Following treatments cells were scanned using a 63X oil immersion lens with the 405 nm laser. Regions of interest (ROI) were created at puncta and fluorescence bleached by rapid scanning of increased laser power (5–10%) to a bleach depth of 40–60%. Time-lapse images were acquired over a 2-min time course post-bleaching at 1-s intervals. Images were processed in Zen and FRAP data were fitted to a single exponential model using GraphPad Prism.

Data analysis was performed using previously published methods.(53) Fluorescence intensities of regions of interest (ROI) in the bleaching area (ROIb = bleached area) were recorded for each time point. The final data was normalized to pre-bleached intensities of the ROIs data and fitted to a single exponential recovery curve. Percent fluorescence recovery (mobile fraction) was calculated from the plateau (Vmax) of the fitted curves normalized to the total bleached fluorescence.

### isoTOP-ABPP sample preparation and LC-MS/MS analysis

Methods for studying changes in cysteine reactivity were adapted from Weerapana, E. *et al*, (15)and Abo, M. *et al*.(54) 1 mg of protein lysates was labeled with either 100 μM IA Alkyne-Light or IA Alkyne-Heavy at room temperature for 1 hr. A photocleavable PC biotin-azide tag was appended to samples by copper-catalyzed azide alkyne cycloaddition (CuAAC) using 10 μM PC biotin azide (Click Chemistry Tools), 1 mM TCEP, 100 μM TBTA, and 1 mM copper (II) sulfate. The reaction was incubated at room temperature for 1 hr with vortexing every 15 min. Samples were combined pairwise and centrifuged at 6,500 g for 10 min at 4 °C. Protein pellets were washed twice in 500 μL ice-cold methanol with gentle sonication and centrifugation at 6,500 g. The supernatant was removed and the protein pellet was re-solubilized in 1 mL 1.2% SDS/DPBS with tip sonication and incubation at 80 °C for 5 min. Protein solutions were combined with 5 mL DPBS and 100 μL streptavidin beads. Samples were incubated at 4 °C overnight. The following day, samples were incubated at room temperature for 2-3 hrs with rotation to re-solubilize SDS and washed with 5 mL of 0.2% SDS/DPBS, 3x 5 mL DPBS, and 3x 5 mL H_2_O. Following each wash, beads were pelleted by centrifugation at 1,400 g for 3 min. The washed beads were re-suspended in 500 μL of 6 M urea with 10 mM DTT and incubated at 65 °C for 20 min with re-suspension every 10 min. 20 mM iodoacetamide was added to the beads and incubated at 37 °C for 30 min with rotation. The reaction was diluted with 950 μL DPBS and centrifuged at 1,400 g for 3 min to remove the supernatant. Beads were re-suspended in 200 μL of 2 M urea, 1 mM CaCl_2_, and 2 μg trypsin. On-bead trypsin digestion was performed overnight at 37 °C with rotation. Beads were washed with 3x 500 μL DPBS and 3x 500 μL H_2_O. Washed beads were re-suspended in 200 μL DPBS and incubated under UV light (365 nm, 80W) for 3 hrs at room temperature. Samples were spun down and the supernatant was collected. Beads were washed twice with 75 μL DPBS and the supernatant was combined with the previous collections to make a final volume of 350 μL. 17.5 μL formic acid was added to the samples. Peptides were packed and de-salted using a 250 μm fused silica desalting column (Agilent) with 4 cm of Aqua C18 reverse phase resin (Phenomenex) using a high-pressure injection cell.

Mass spectrometry data was obtained using a Thermo Fisher LTQ Orbitrap Discovery coupled to an Agilent 1200 series HPLC. Peptides were eluted onto a 100 μm fused silica biphasic column with a 5 μm tip, packed with 10 cm C18 resin and 4 cm of Partisphere strong cation exchange (SCX) resin. The gradient for peptide elution and separation ranged from 0-100% buffer B (80% acetonitrile, 20% H_2_O, 0.1% formic acid) in buffer A (95% H_2_O, 5% acetonitrile, 0.1% formic acid). Peptides were eluted from SCX to C18 using 4 separate salt pushes (50%, 80%, 100%, 100%) followed by a gradient of buffer B in buffer A as according to Weerapana, E. *et al*(*48*) The flow rate was set to 0.25 μL/min and the spray voltage was set to 2.75 kV. One full MS scan was followed by 8 data-dependent scans of the n^th^ most intense ions. Dynamic exclusion was enabled for each run.

MS data was searched using the SEQUEST algorithm (55) against a concatenated target/decoy non-redundant variant of the *Mus Musculus* FASTA UniProt database.(49) A static modification of +57.02146 on cysteine was included for cysteine alkylation by iodoacetamide. Differential modifications of +228.13749 (light) and +234.15762 (heavy) on cysteine were included to account for IA-Light or IA-Heavy appended to a cleaved PC biotin-azide tag. SEQUEST output files were filtered using DTASelect 2.0(*50*) with -trypstat and -modstat options and a maximum false discovery rate (FDR) of 5% was applied. Peptides were required to be fully tryptic (-y 2), with a found modification (-m 0), a delta-CN score greater than 0.06 (-d 0.06), and at least one peptide was required per locus (-p 1). H:L ratios were quantified using CIMAGE quantification package.

### OxiCAT sample preparation and LC-MS/MS analysis

OxICAT sample preparations were adapted from Bechtel, T. *et al*,(27) and Topf, U. *et al*.(56) 100 μg of cell lysates was precipitated in 100% TCA/DPBS at -80 °C for 1 hr or overnight. Samples were pelleted at 14,000 rpm for 10 min at 4 °C and resuspended in 500 μL of ice-cold acetone by vortexing. Proteins were pelleted at 5,000 rpm for 10 min at 4 °C. Acetone was removed and the pellet was left air dry to remove traces of acetone. The pellet was resuspended in 100 μL of DAB buffer (6 M urea, 20 mM Tris-HCl, 1 mM EDTA, 0.5% SDS, pH 8.5) and incubated with 10 mM N-ethylmaleimide (NEM) for 2 hrs at 37 °C with agitation. The reaction was first diluted 3-fold with H_2_O and then 5-fold with ice-cold acetone. Proteins were precipitated at -20 °C for 2 hrs or overnight and pelleted at 4,500 g for 30 min at 4 °C. The protein pellet was re-suspended in 500 μL of ice-cold acetone and centrifuged at 4,500 g for 10 min. Proteins were re-solubilized using 80 μL DAB buffer and incubated with 2.5 mM TCEP for 5 min at 37 °C. The reaction was further diluted with 120 μL DAB buffer and incubated with 10 mM D5-NEM (Cambridge Isotope Laboratories) for 2 hrs at 37 °C. The reaction was diluted 3-fold with H_2_O and then 5-fold with ice-cold acetone for protein precipitation at -20 °C for 2 hrs or overnight. Proteins were pelleted at 4,500 g for 30 min at 4 °C and washed with 500 μL ice-cold acetone. Protein pellets were re-suspended in 200 μL 2 M urea with 1 mM CaCl_2_ and 2 μg trypsin for overnight digestion at 37 °C. 10 μL formic acid was added to the digested peptides. Samples were de-salted using Sep-Pak C18 columns and eluted with 1.5 mL of 80% LC/MS-grade acetonitrile, 20% H_2_O, and 0.1% formic acid. The de-salted samples were dried on SpeedVac for further storage at -20 °C.

Trypsin digested peptide samples were resuspended in 100 μL buffer A peptide concentration was determined using Pierce^TM^ quantitative colorimetric peptide assay. Peptide analysis was performed using an Orbitrap Exploris 240 mass spectrometer coupled to a Dionex Ultimate 3000 RSLCnano system. 500 ng peptides were loaded onto an Acclaim PepMap 100 C18 loading column. Peptides were eluted onto an Acclaim PepMap RSLC column and separated with a 160-min gradient ranging of 5-25% buffer B in buffer A at a flow rate of 0.3 μL/min. The spray voltage was set to 2.1 kV. One full MS scan (350-1,800 m/z, 120,000 resolution, RF lens 65%, automatic gain control (AGC) target 300%, automatic maximum injection mtime, profile mode) was obtained every 2 s with dynamic exclusion (repeat count 2, duration 10 s), isotopic exclusion (assigned), and apex detection (30% desired apex window) enabled. A variable number of MS2 scans (15,000 resolution, AGC 75%, maximum injection time 100 ms, centroid mode was obtained between each MS1 scan based on the highest precursor masses, filtered for monoisotopic peak determination, theoretical precursor isotopic envelope fit, intensity (5E4), and charge state (2–6). MS2 analysis consisted of the isolation of precursor ions (isolation window 2 m/z) followed by higher energy collision dissociation (HCD) (collision energy 30%).

Analysis of MS data collected on Orbitrap Exploris 240 was performed using Thermo Proteome Discoverer software. Protein identification was achieved using the SequestHT and Percolator algorithms(57) against *Mus Musculus* proteome UniprotKB database.(49) The protease enzyme was set as trypsin with a maximum of 2 missed cleavages allowed. The peptide precursor mass tolerance was set to 10 ppm with a fragment mass tolerance of 0.02 Da. Dynamic modifications included methionine oxidation (+15.995), N-terminal acetylation (+42.011), methionine loss (-131.040), and cysteine alkylation by either light-NEM (+125.048) or heavy-NEM (+130.079). The FDR for highly confident peptide identification was set at 1%. The resulting H:L ratios were converted to percent oxidation values by the following equation: 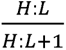. Average percent oxidation values were calculated for each peptide and peptides with standard deviation >50% were removed from the analyses.

#### Human intestinal cells

Healthy human duodenal cells were generated from biopsies obtained from a patient during diagnostic endoscopy as part of routine clinical care under approved Boston Children’s Hospital Institutional Review Board Protocol # P00027983.

#### Antibodies

RACK1 (CST 5432S), β-actin (Fisher Scientific A5441), p65 (CST, 8242S), IRF3 (CST, 11904S),Flag (F7425 Sigma), HA (CST 3724S), Puromycin (Sigma MABE343-AF647), pNFKB (CST 3033S),IKKA (CST 61294S), G3BP1(Abcam, 56574), p38 MAPK(CST 8690T),: p65 (CST, 8242S) and IRF3 (CST, 11904S).

#### Cell Line

HEK293T (ATCC), HCT8 cells (ATCC)

#### Mouse

WT: C57BL/6J Strain Jackson Lab #:000664 NOX1KO: B6.129X1-Nox1tm1Kkr/J Strain Jackson Lab #: 018787

## Extended Data

**Figure S1.**
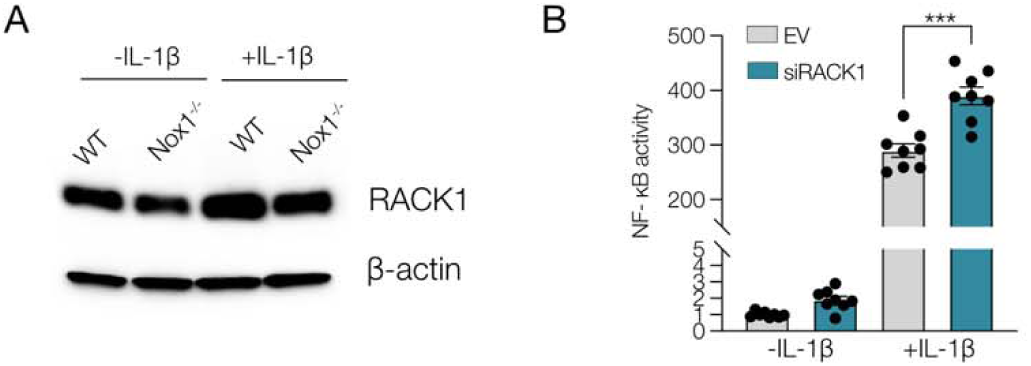
**(A)** Biotin switch immunoprecipitation and immunoblot with anti-RACK1 showing increased proportion of oxidized RACK1 following IL-1β (10 ng/μL) in WT versus NOX1 knockout colonic epithelial cells. **(B**) NF-κB activity in HEK cells expressing scrambled vector (EV), or knockdown of RACK1 (siRNA) following IL-1β (10 ng/μL). (n≥8) ***p < 0.001

**Figure S2.**
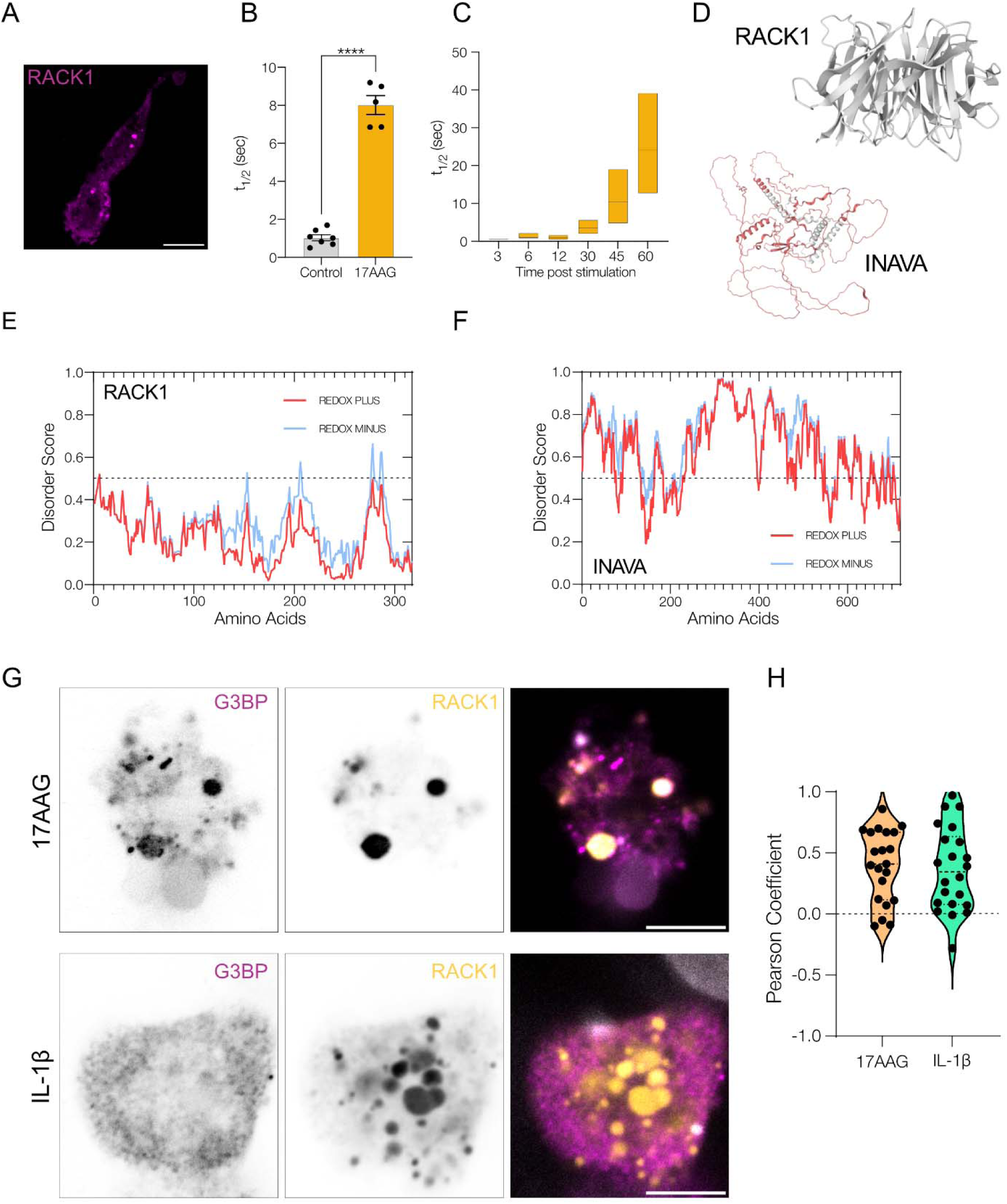
**(A)** Representative example of HCT-8 transfected with RACK1-mkate forming puncta following treatment with H_2_O_2_. **(B)** Diffusion properties of RACK1 puncta following stimulation with 17-AAG (10 µM) relative to control. (n ≥2) **(C)** Diffusion properties of RACK1 puncta at different times following stimulation with 17-AAG (10 µM). (n =2) **(D)** PDB structure of RACK1 with almost no unstructured regions and PDB structure of INAVA as predicted by Alphafold showing large regions of highly unstructured regions and low -residue model confidence score (shaded in red). **(E)** Predicted Disorder score of RACK1 with and without changes in redox state of amino acids in protein. The dotted line at 0.5 represents the cut-off value for disordered region. **(F)** Predicted Disorder score of INAVA with and without changes in redox state of amino acids in protein. **(G)** Representative immunofluorescence image of HCT-8 overexpressing wt RACK1-mcherry and stained with G3BP and stimulated with 17-AAG (10 µM) IL-1β (10 ng/μL). Top is a representative example with a cell with high colocalization and bottom represents a cell with low colocalization. **(H)** Pearson correlation coefficient showing colocalization of G3BP and RACK1 following stimulation with either 17-AAG (10 µM) or IL-1β (10 ng/μL). (G) was obtained from three independent biological repeats.

**Figure S3.**
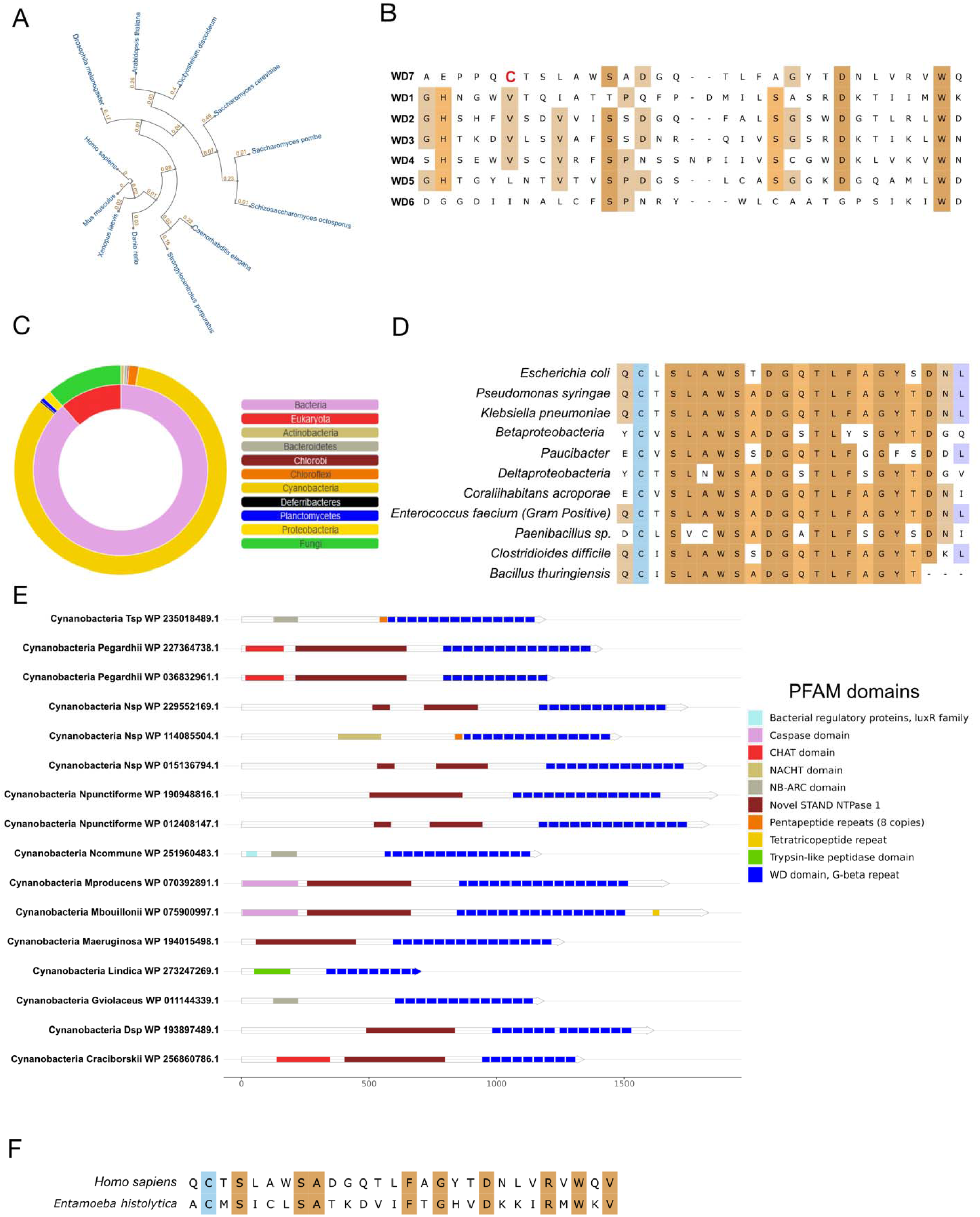
**(A)** Evolutionary distance of different eukaryotic RACK1 **(B)** Sequence alignment of 7WD domains of human RACK1. **(C)** RACK1 homologs across diverse kingdoms as a sunburst plot. **(D)** Sequence alignment of different bacterial homologs **(E)** PFAM analysis of homologs of RACK1 in cyanobacteria showing diverse combinatorial domains in proteins with WD domains. **(F)** WD7 sequence alignment between human and Entamoeba *histolytica* RACK1 showing conserved cysteine.

**Figure S4.**
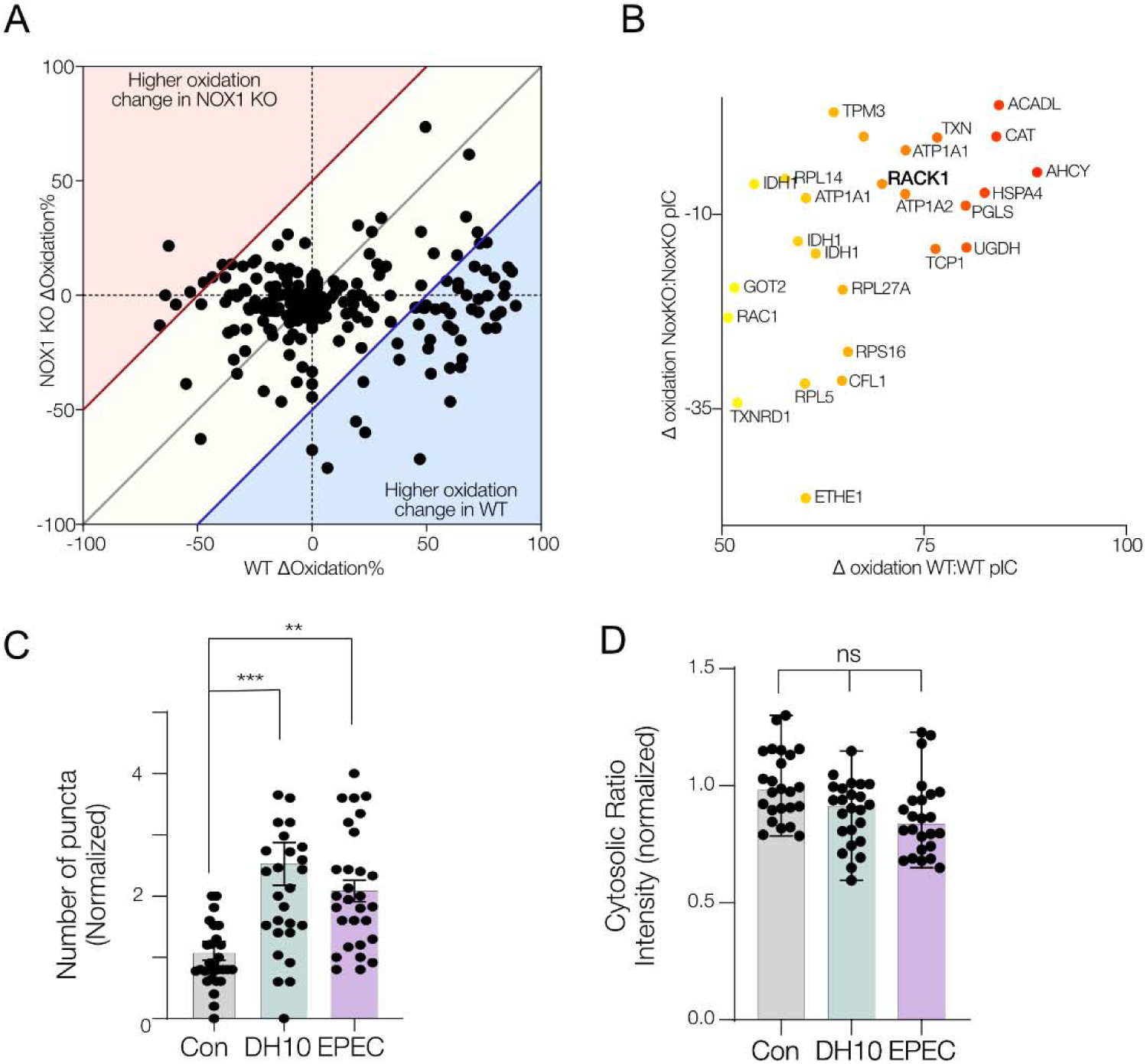
**(A)** Scatter plot showing the percent oxidation of cysteine residues following stimulation by polyI:C (10ug/ml) by OxiCAT mass spectrometry of proteins in NOX1^-/-^ and WT organoids. **(B)** Subset of proteins with cysteines significantly increased oxidation in WT and low or reduced oxidation in NOX1^-/-^ following polyI:C stimulation. **(E)** Quantification of number of puncta formed per transfected cell. **(F)** Quantification of number of Mean cytosolic intensity ratio (405/488) per transfected cell. (E) and (F) were obtained from three independent biological repeats. *p < 0.05, **p < 0.01, ***p < 0.001

