## Supplementary Figures for "Trans-kingdom coupling of redox signaling to environmental cell stress responses through multiphase partitioning"

### Extended Data

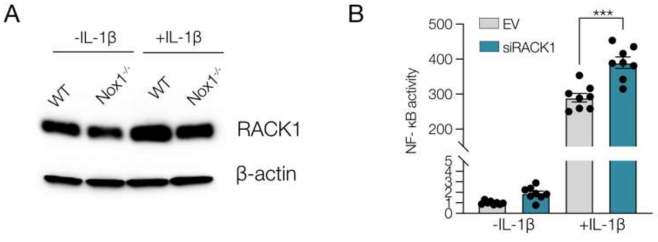


Figure S1

**Figure S1 (A)** Biotin switch immunoprecipitation and immunoblot with anti-RACK1 showing increased proportion of oxidized RACK1 following IL-1β (10 ng/µL) in WT versus NOX1 knockout colonic epithelial cells. **(B)** NF-κB activity in HEK cells expressing scrambled vector (EV), or knockdown of RACK1 (siRNA) following IL-1β (10 ng/µL). (n≥8) ***$p < 0.001$

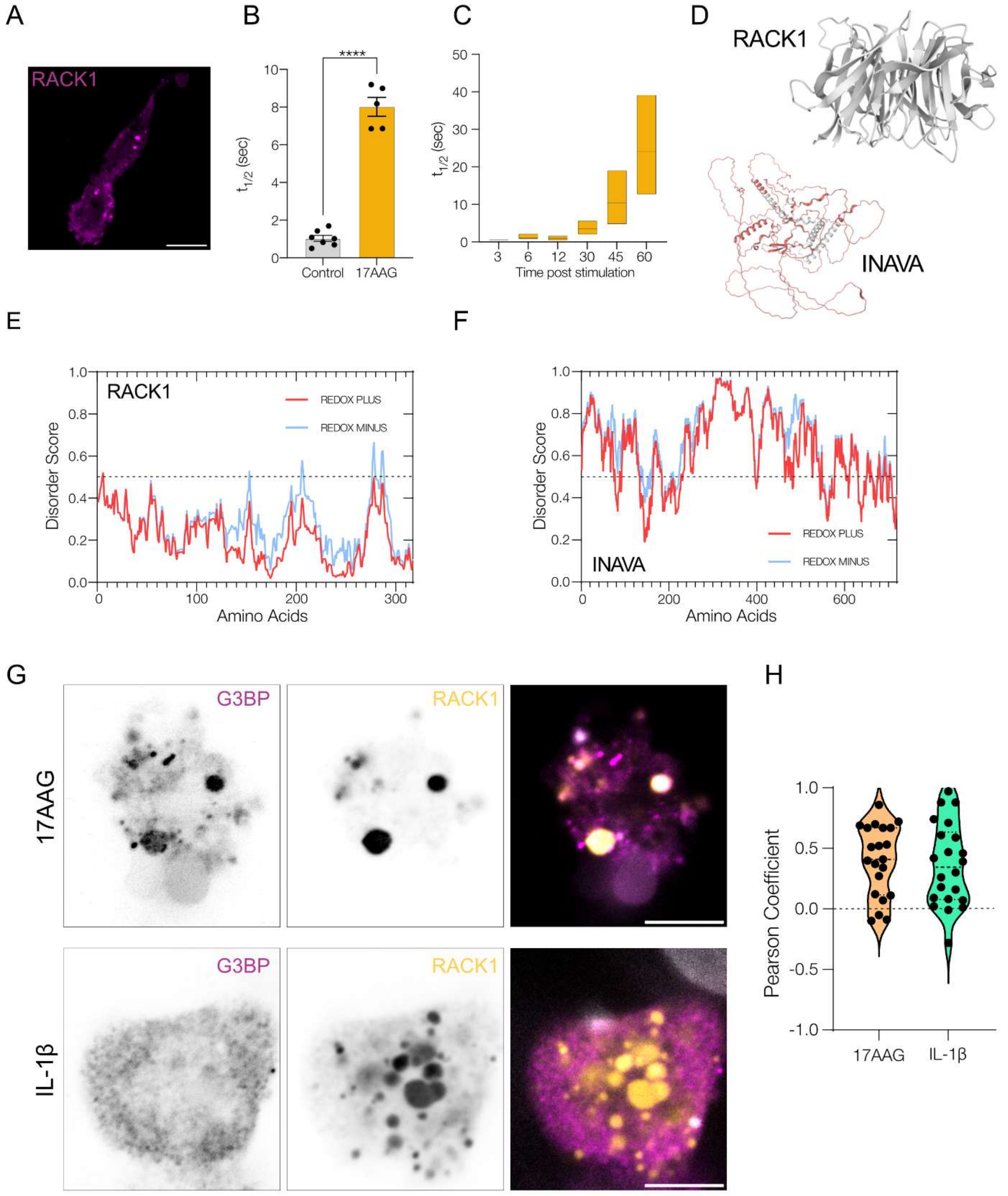


**Figure S2 (A)** Representative example of HCT-8 transfected with RACK1-mkate forming puncta following treatment with $H_2O_2$. **(B)** Diffusion properties of RACK1 puncta following stimulation with 17-AAG (10 µM) relative to control. (n ≥2) **(C)** Diffusion properties of RACK1 puncta at different times following stimulation with 17-AAG (10 µM). (n =2) **(D)** PDB structure of RACK1 with almost no unstructured regions and PDB structure of INAVA as predicted by Alphafold showing large regions of highly unstructured regions and low -residue model confidence score (shaded in red). **(E)** Predicted Disorder score of RACK1 with and without changes in redox state of amino acids in protein. The dotted line at 0.5 represents the cut-off value for disordered region. **(F)** Predicted Disorder score of INAVA with and without changes in redox state of amino acids in protein. **(G)** Representative immunofluorescence image of HCT-8 overexpressing wt RACK1-mcherry and stained with G3BP and stimulated with 17-AAG (10 μM) IL-1β (10 ng/μL). Top is a representative example with a cell with high colocalization and bottom represents a cell with low colocalization. **(H)** Pearson correlation coefficient showing colocalization of G3BP and RACK1 following stimulation with either 17-AAG (10 μM) or IL-1β (10 ng/μL). (G) was obtained from three independent biological repeats.

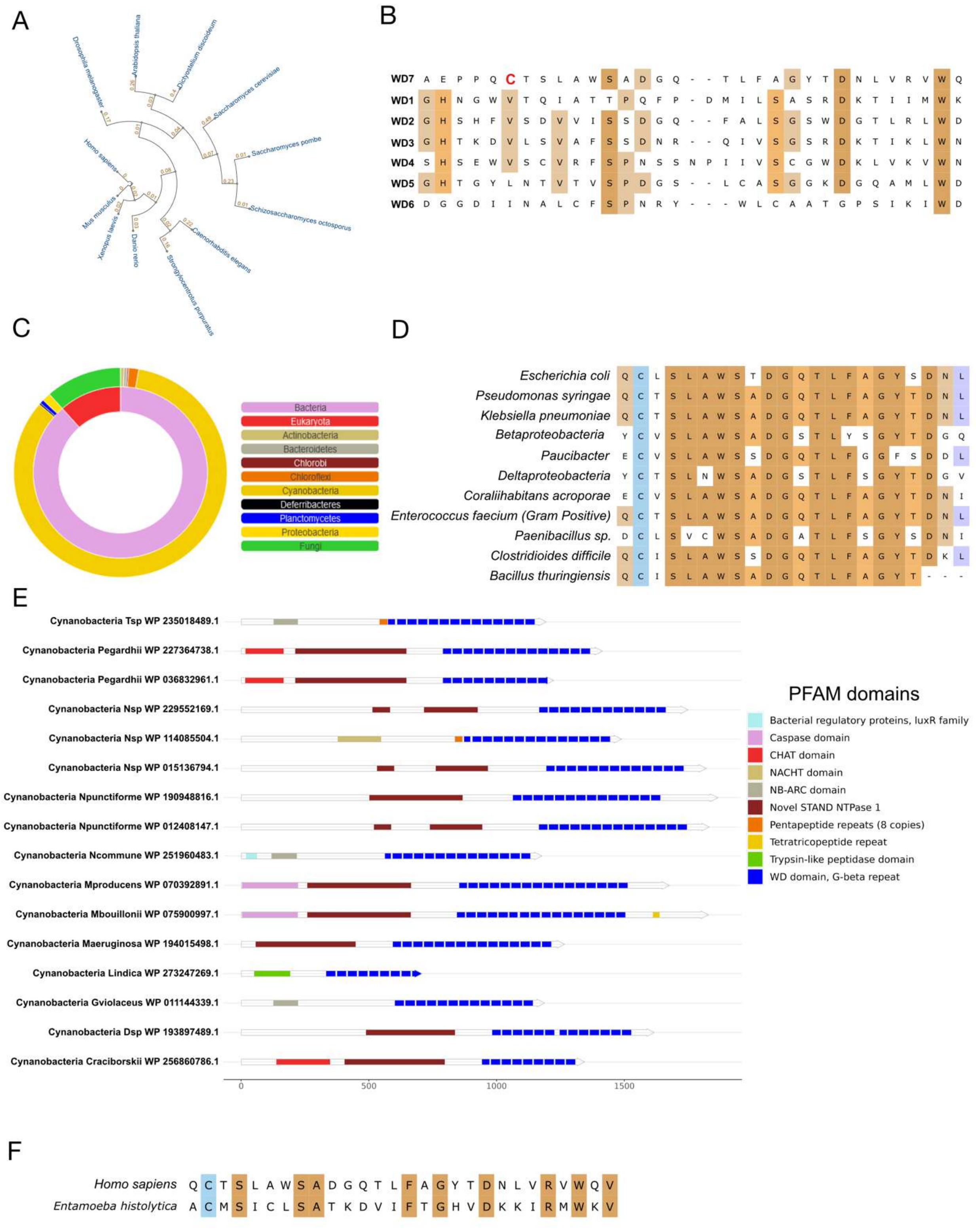


Figure S3

**Figure S3 (A)** Evolutionary distance of different eukaryotic RACK1 **(B)** Sequence alignment of 7WD domains of human RACK1. **(C)** RACK1 homologs across diverse kingdoms as a sunburst plot. **(D)** Sequence alignment of different bacterial homologs **(E)** PFAM analysis of homologs of RACK1 in cyanobacteria showing diverse combinatorial domains in proteins with WD domains. **(F)** WD7 sequence alignment between human and Entamoeba *histolytica* RACK1 showing conserved cysteine.

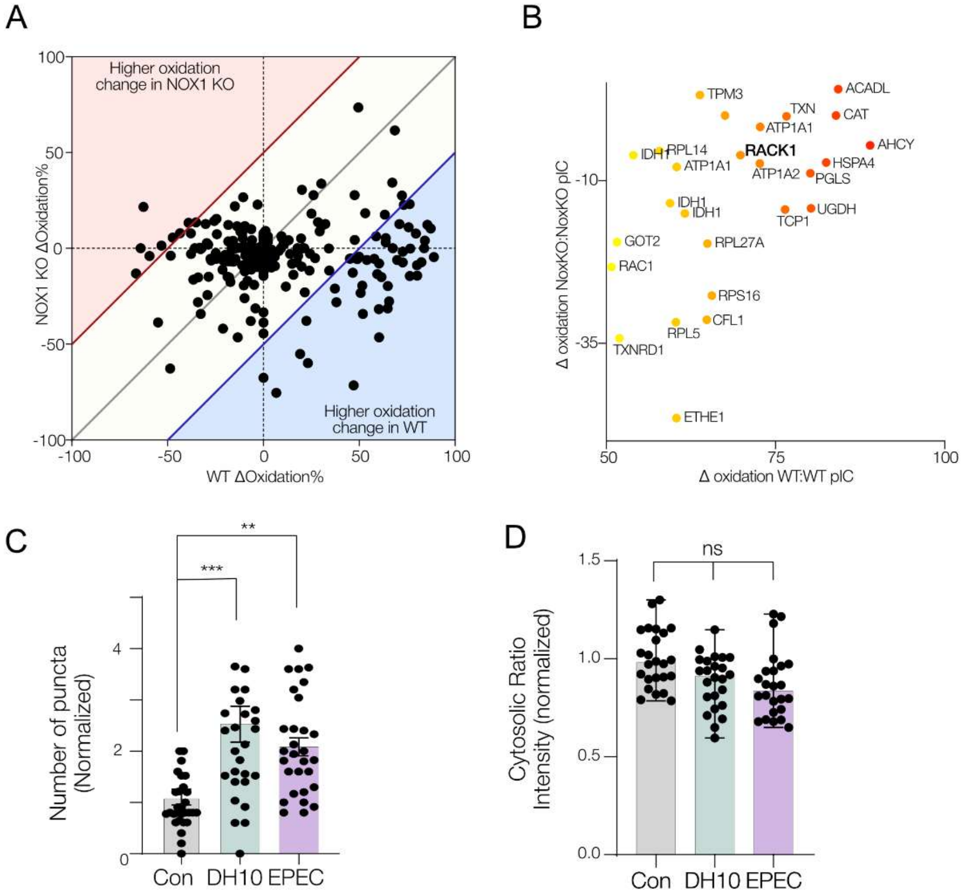


**Figure S4 (A)** Scatter plot showing the percent oxidation of cysteine residues following stimulation by polyI:C (10ug/ml) by OxiCAT mass spectrometry of proteins in NOX1$^{-/-}$ and WT organoids. **(B)** Subset of proteins with cysteines significantly increased oxidation in WT and low or reduced oxidation in NOX1$^{-/-}$ following polyI:C stimulation. **(E)** Quantification of number of puncta formed per transfected cell. **(F)** Quantification of number of Mean cytosolic intensity ratio (405/488) per transfected cell. (E) and (F) were obtained from three independent biological repeats. *p < 0.05, **p < 0.01, ***p < 0.001
